# Hepadnavirus tropism determined by diversified NTCP functionality in Feline species

**DOI:** 10.64898/2026.11.10.765656

**Authors:** Junki Mifune, Toru Ekimoto, Junko S Takeuchi, Chisa Kobayashi, Misaki Kurata, Kaho Shionoya, Miki X Kawano, Akatsuki Saito, Camille Sureau, Mitsunori Ikeguchi, Koichi Watashi

## Abstract

Hepatitis B virus (HBV) exhibits a narrow host range, largely determined by sodium taurocholate cotransporting polypeptides (NTCP), a hepatic bile acid transporter acting as the HBV receptor. NTCP among animals appears to have evolved following trade-off mutations to lose its viral receptor activity while preserving its transporter function. Here, through profiling of the NTCP homologs from 23 mammalian species, we uncover unexpected functional heterogeneity of feline NTCPs. NTCPs derived from cat, tiger, and leopard supported viral preS1 binding and HBV infection, whereas lion-NTCP was deficient for preS1 binding and, notably, exhibited markedly reduced bile acid transport. Cat-NTCP was further demonstrated to have viral receptor function by productive infection in primary cat hepatocytes. Experimental mutagenesis identified double substitution at positions 191 and 268 as key determinants: introducing lion-type substitutions into cat-NTCP (G191R and T268A) abolished preS1 binding, whereas reciprocal substitutions in lion-NTCP restored receptor function. Molecular dynamics simulations suggested that lion-NTCP adopts conformation in which a transmembrane helix bending enlarges the whole bile acid tunnel and disrupts preS1 compatibility. Such differences also determined host susceptibility to another feline-infected domestic cat HBV. These findings reveal diversified NTCP functionality with evolutionary plasticity that acquires resistance to hepadnavirus infection at the expense of physiological transporter function.

## Introduction

Hepatitis B virus (HBV), a member of the *Hepadnaviridae* family, infects a limited range of host species, primarily hominoids such as humans and chimpanzees. Despite a phylogenetically close lineage to humans, HBV does not infect Old World Monkeys (OWM) such as cynomolgus macaques[1–6]. However, the full spectrum of HBV-susceptible animals is not precisely defined. Its closely related orthohepadnaviruses that infect diverse mammalian hosts, including New World Monkeys (NWM) (e.g., woolly monkeys and capuchin monkeys), domestic cats and dogs, horses, several bat species, and rodents such as squirrels and woodchucks have been reported[7–17]. These orthohepadnaviruses are reported to infect specific species, yet their host ranges and determinants for cross-species transmission are poorly analyzed.

HBV infection to host cells is primarily mediated by the N-terminal region of the large HBV surface antigen (LHBs), known as the preS1 region, which is displayed at the outermost surface of the virion. The N-terminally myristoylated 2–48 amino acids of preS1 (myr-2–48^preS1^) is essential for HBV binding to its host receptor, sodium taurocholate cotransporting polypeptide (NTCP), a liver-specific bile acid transporter[18,19]. Through interaction with NTCP on cell surface, HBV enters into hepatocytes and delivers its relaxed circular DNA genome to the nucleus, where it is converted into covalently closed circular DNA (cccDNA), a template for genome replication and virus production[20]. HBV’s narrow host range is largely due to incapability of viral entry in most non-hominoid animal species: while hepatocytes derived from most mammalians can support HBV genome replication from cccDNA and the following virus particle production, only a limited number of species allows productive HBV entry[21,22]. Restriction during viral entry process is likely to involve multiple mechanisms, depending on species, a typical mechanism of which is loss of virus receptor functionality of the NTCP homologs (see below). The NTCP-mediated entry pathway, and the resulting narrow host tropism, applies also to hepatitis D virus (HDV), a satellite virus that co-infects with HBV and carries HBV surface antigens on its virion surface, which therefore relies on NTCP for cell entry and is frequently used as an experimental model for analyzing the early steps of viral entry.

Recently, we and other groups have solved the cryo-electron microscopy (cryoEM) structures of human NTCP and its complex with myr-2–48^preS1^ [23–28]: NTCP is a nine transmembrane (TM) protein composed of a panel domain (with TM1, 5, and 6) and a core domain (with TM2, 3, 4, 7, 8, and 9), together forming a bile acid transporter tunnel between the two domains[24]. This overall architecture is conserved among all the reported cryoEM structures of NTCP from mammalian species, including human, macaque, cow, and rat[23,29]. Binding of preS1 relies on the large area of the NTCP structure, ranging from the bile acid tunnel to the outersurface loops (see Fig. 8)[26,27,30]. Species-specific sequence substitutions in these NTCP areas can modify preS1 compatibility that leads to escape from infection. However, because of a highly structured nine-TM protein in a relatively short sequence (349 aa in human), NTCP contains only limited regions for sequence diversity with securing the original bile acid transporter function. Two short amino acid motifs, 84–87 aa (Species Determinant Motif-1, SDM-1) and 157–165 aa (SDM-2), are known as mutation hot spots among species for determining infection susceptibility[31] : The 157–165 aa region in TM5 demernines the preS1 compatibility on the bile acid tunnel, of which sequence variation in the OWM is responsible for their resistance to HBV/HDV infection (see Fig. 8-ii)[5,6,18]. The substituted sequence at the extracellular 84–87 aa region in the TM2-TM3 outer-loop leads to resistance to HDV in murine and low HBV susceptibility in NWM (see Fig. 8-iii)[3,29,32–34]. These two motifs have evolved under positive selection, likely as a result of selective pressure by hepadnavirus infection[6,35]. Sequence variation in these regions affects only the local structure but retains the overall protein architecture, securing the original bile acid transporter function[23,29]. Thus, heterogeneity of NTCP function among species reported to date appears to result from the trade-off between escaping infection and preserving transporter function. However, in this study, we unexpectedly found that NTCPs derived from feline species had diverse virus receptor function (see Figs 1–2), despite their complete sequence conservation in the above two motifs. Analysis for these differences could disclose a yet unidentified mechanism by which NTCP heterogeneity governs hepadnavirus susceptibility and constitutes a barrier against interspecies transmission.

**Fig. 1.**
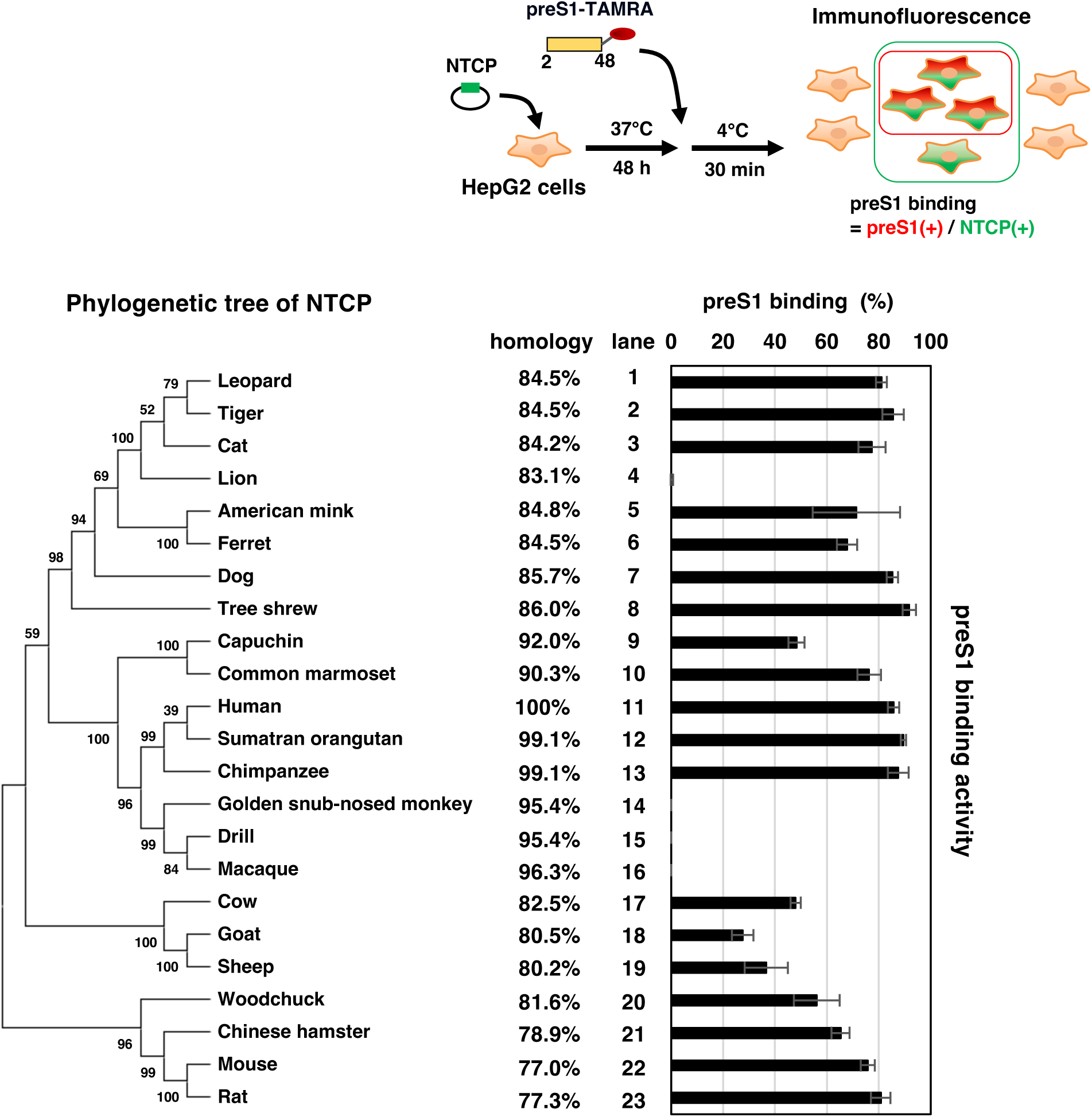
Capacity of mammalian NTCPs to support preS1 binding. Upper, schematic representation of preS1 binding assay. HepG2 cells were transfected with the expression plasmid expressing the myc-tagged NTCP derived from each mammalian and were incubated with 250 nM myristoylated preS1 peptide, consisting of 2–48 aa region conjugated with TAMRA (preS1-TAMRA). Cell-bound TAMRA fluorescence as well as NTCP detected with anti-myc antibody by immunofluorescence analysis were visualized with a fluorescence microscopy. Lower left, phylogenetic tree of NTCP derived from 23 mammalian species built using the Neighbor-Joining method. Lower right, the numbers of TAMRA- and NTCP-positive cells were counted and the percentages of preS1-positive cells over NTCP-positive cells are shown in the graph.

**Fig. 2.**
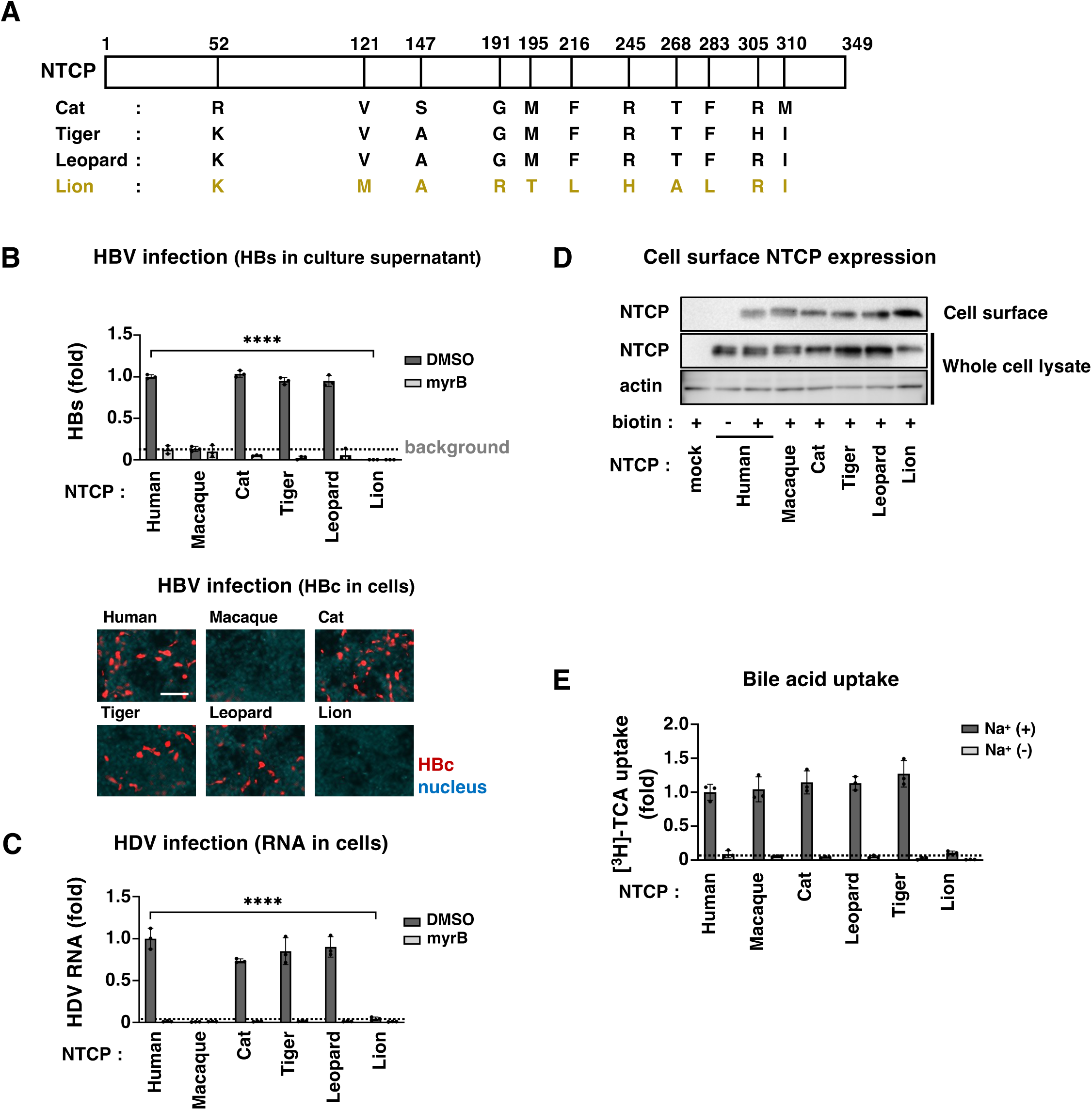
Capacity of viral receptor function of feline family-derived NTCPs. **(A)** Amino acids comparison between NTCPs derived from feline families (cat, tiger, leopard, and lion). Non-conserved amino acids and their amino acid numbers in NTCPs are indicated. **(B)** HBV infection assay. HepG2 cells transfected with the expression plasmid encoding each NTCP were inoculated with HBV for 16 h either in the presence (light gray) or the absence (dark gray) of myrcludex-B (myrB), a positive control to inhibit HBV infection. HBV infection was evaluated by quantifying HBs antigen in the culture supernatant at 12 days post-inoculation by ELISA (upper) and by detecting intracellular HBc antigen by Immunofluorescence (lower). The dotted line in the upper graph indicates the background levels. Red and blue signals in the lower pictures indicate HBc and nucleus, respectively. The scale bar in the pictures, 100 µm. **(C)** HDV infection assay. The cells transfected as shown in (b) were inoculated with HDV for 16 h either in the presence (light gray) or the absence (dark gray) of myrB. HDV infection was evaluated by quantifying intracellular HDV RNA at six days post-inoculation by real time RT-PCR. **(D)** Expression levels of NTCP both on cell surface (upper) and in total cell (middle) as well as actin in total cell as an internal control (lower) were detected by immunoblot. Cell surface protein fraction was recovered by treating the cells with (+) or without (-) the biotinylated solution that is not cell membrane permeable, followed by lyzing cells and pull down with streptavidin-beads as shown in Methods. **(E)** Bile acid uptake activity of feline family-derived NTCPs were measured by incubating the NTCP-transfected cells as shown in (b) with [^3^H]-taurocholic acid as a substrate either upon sodium-free (light gray) or -containing buffer (dark gray) for 10 min and counting the intracellular radioactivity with liquid scintillation counter.

In this study, from the functional profiling of NTCPs derived from 23 mammalian species, we showed that NTCPs derived from cat, tiger, and leopard, supported HBV/HDV infection, whereas lion-derived NTCP did not. This functional difference was determined by combined amino acid substitutions, but not a single, at aa 191 (in TM6) and 268 (in TM8b) on the bile acid tunnel. Unexpectedly, lion NTCP showed marked impairment for both bile acid transport and viral receptor functions, due to a potential distinct structure in the bile acid tunnel. These NTCP variations also governed the susceptibility to domestic cat HBV, originally infected in domestic cats. Our finding reveals a yet undefined diversity of NTCP transporter function that constitutes a barrier against interspecies transmission of hepadnavirus.

## Results

### Capacity of mammalian-derived NTCPs to support preS1 binding

To profile the function of mammalian NTCPs for supporting preS1 binding, we selected 23 NTCPs derived from mammalian species including hominoids, OWM and NWM, *Canidae*, *Mustelidae*, *Felidae*, *Bovidae*, and rodents, as well as tree shrew, of which sequence is registered in the public database (Table S1). We built a phylogenetic tree of these NTCPs, and showed their amino acid sequence homologies to human NTCP (Fig. 1): Primate NTCPs form one cluster (Fig. 1, lanes 9–16), with its closest cluster including cat, dog, and tree shrew (Fig. 1, lanes 1–8), and a distant cluster including rat and mouse (Fig. 1, lanes 22 and 23). We constructed the expression plasmids for these 23 NTCPs and transfected them into HepG2 cells (originally deficient for NTCP expression), which were incubated with TAMRA fluorescence-conjugated preS1 peptide consisting of 2–48 aa region with N-terminal myristoylation (myr-2–48^preS1^-TAMRA)[26,36], to evaluate the NTCP’s capacity to support preS1-mediated binding (Fig. 1, upper). To normalize the variation of NTCP overexpression levels, we detected NTCP as well as cell-bound TAMRA fluorescence signals by immunofluorescence and calculated the signal area of preS1-TAMRA signals divided by that of NTCP signals as percentage (Figs 1 and S1 Fig). As shown in Supplementary Fig. 1, NTCP derived from hominoids such as Human, Chimpanzee, and Sumatran orangutan, showed preS1 binding in 85–90% of cells (Figs 1, lanes 11-13 and S1 Fig), whereas that of OWM (Golden snub-nosed monkey, Dill, and Macaque) supported no or negligible preS1 binding (Figs. 1, lanes 14–16 and S1 Fig), consistent with the accumulated reports showing that hominoids but not Old World Monkeys are susceptible to HBV/HDV infection[5,6]. Also, as reported previously[3,5,18,19,32,37], NTCPs derived from cat, dog, tree shrew, common marmoset, mouse, and rat bound to preS1, with varying binding levels at 71–91% (Figs 1, lanes 3, 7, 8, 10, 22, and 23 and S1 Fig). Among the overall preS1 binding profile, feline familly has a clear contrast in which NTCP from leopard, tiger, and cat suported preS1 binding at the equivalent levels to that for hominoid NTCPs, but lion-derived NTCP showed little preS1 binding (Figs 1, lanes 1–4 and S1 Fig). We then focused on these feline NTCPs as a model for analyzing the diverse activity to determine virus susceptibility, despite within a genetically close cluster.

### Cat-, tiger-, and leopard-derived NTCPs are functional for both virus receptor and bile acid transpoprter, but lion NTCP does not

Feline familly-derived NTCPs have 97% indentity in amino acid sequence, with 11 different residues (aa 52, 121, 147, 191, 195, 216, 245, 268, 283, 305, 310) out of 349 amino acids (Fig. 2A). We examined the functionality of these NTCPs as a viral receptor in HBV infection assay, where HepG2 cells were overexpressed with these NTCPs and inoculated with HBV to examine the HBs and HBc productions as HBV infection markers at 12 days post-inoculation. As shown in Fig. 2B, HepG2 cells overexpressing NTCPs derived from cat, tiger, and leopard showed the production of HBs in the culture supernatant and of HBc in the cells at the levels equivalent to human NTCP-overexpressing cells, and which were lost by treatment with an HBV/HDV entry inhibitor, myrcludex-B (myrB) [6,18,29] (Fig. 2B). In contrast, lion NTCP-expressing cells produced almost no HBs and HBc upon HBV inoculation, around or below the levels for cells overexpressing NTCP derived from macaque, a HBV/HDV non-susceptible species[6,29] (Fig. 2B). Similar results were obtained in HDV infection assay (Fig. 2C), consisitent with HBV and HDV sharing the same early entry mechanism[18,19]. These results were not due to the altered expression or subcellular localization of NTCP itself, as shown by the equivalent expression levels for NTCPs in cell surface (Fig. 2D) as well as the dominant cell surface NTCP distribution by confocal microscopy immunofluorescence analysis (S2A Fig), for all these NTCPs from different species. We also examined the bile acid uptake activity of these NTCPs by transporter assay, in which HepG2 cells were overexpressed with these NTCPs and were incubated with [^3^H]-taurocholic acid either in the presence or absence of Na^+^ for 10 min to measure the intracellular radioactivity. Bile acid uptake inside cells in the presence of Na^+^ indicates the Na^+^-dependent NTCP transporter acitivity[20]. As shown in Fig. 2E, cat, tiger, and leopard NTCP showed Na^+^-dependent bile acid transport as for human or macaque NTCP, but surprisingly, lion NTCP showed almost negligible bile acid uptake near the background level (Fig. 2E). To assesse whether lion-derived NTCP exhibits low-level bile acid uptake activity or completely lacks transporter capability, we examined transporter kinetics analysis in human, cat, and lion NTCP expressing cells, as reported[38]. As shown in S2B Fig, [^3^H]-TCA uptake was observed in lion NTCP-expressing cells in a substrate concentration-depending manner, but its kinetics profile was different from that of human- and cat-derived NTCPs (S2B Fig). The maximum amount of [^3^H]-TCA uptake, *V_max_* normalized by the expression levels of cell surface NTCP protein, was calculated to be 5327, 6869, and 922 for human, cat, and lion NTCP, respectively, indicating that the maximum bile acid uptake by lion NTCP is only 13–18% of that by human and cat NTCP. The above data suggest that cat, tiger, and leopard-derived NTCPs possess both the viral receptor and bile acid transporter functions, whereas lion-derived NTCP was deficient as a viral receptor and seriously low bile acid transporter function.

We further examined whether cat-derived hepatocytes are really susceptible to infection by using primary cat hepatocytes (PCH) as well as primary rat hepatocytes (PRH) as a negative control. These cells were inoculated with HDV at varying amounts, followed by detection of intracellular HDV RNA and HDAg at six days post-inoculation. As shown in Fig. 3, intracellular HDV RNA was detected in PCH along with the increasing amount of HDV inoculum, and these signals were disappeared to around the background level upon treatment with myrB, suggesting that the detected HDV RNA was derived from HDV infection (Fig. 3). No such signals were detected in PRH with any inoculated HDV amounts, as expected (Fig. 3). Consistently, HDAg was clearly detected with increasing amounts of inoculated virus in PCH, but not PRH, and all the HDAg detection was lost by treatment with myrB (Fig. 3). We tried HBV infection to these hepatocytes, but it was not technically successful because of unhealthy culture condition of these PCH up to two week culture, which is required in a standard HBV infection assay. However, these results at least suggest that NTCP expressed in primary cultures of cat hepatocytes is functional for supporting infection. It was very recently reported that PCH could be infected with HBV as well as HDV[39], consistent with our conclusion that cat NTCP supports the receptor function for HBV and HDV.

**Fig. 3.**
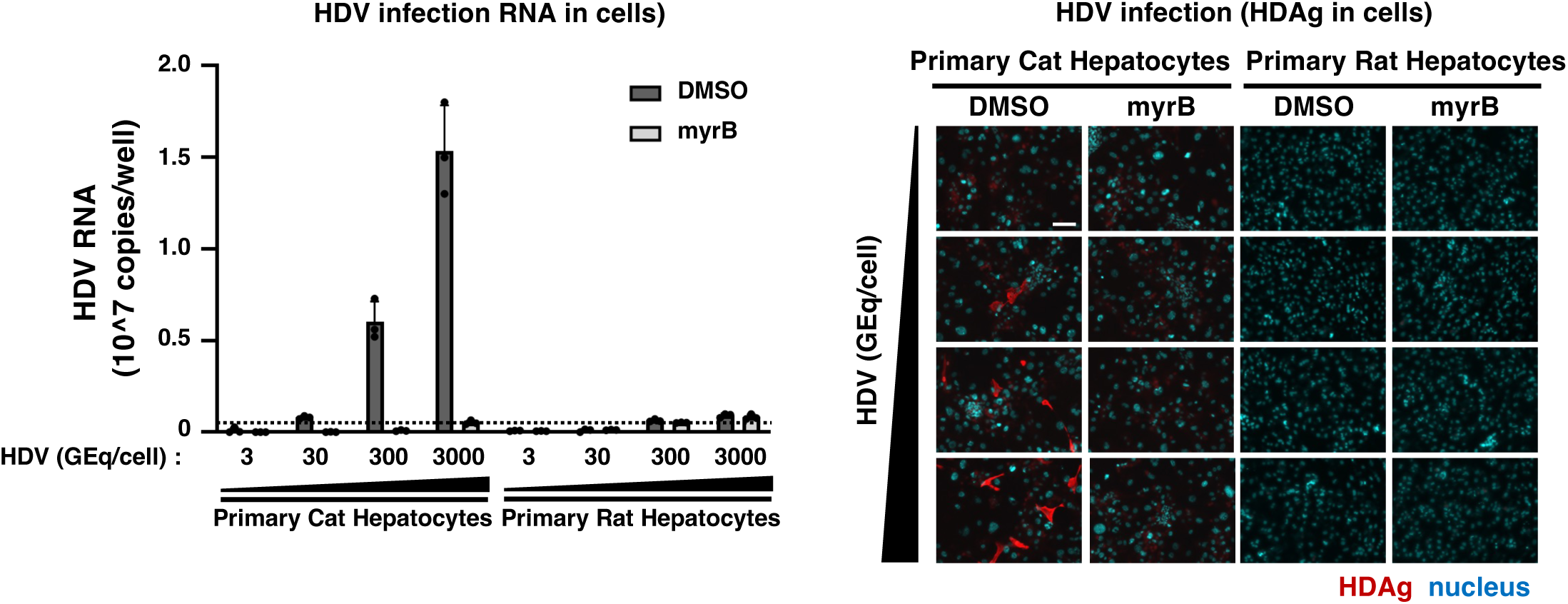
Primary cat hepatocytes were susceptible to HDV infection. Primary hepatocyte cultures derived from cat and rat, as a negative control, were used for HDV infection assay with a various amount of HDV inocula (3, 30, 300, and 3,000 GEq/cell) in the presence (light gray) or absence (dark gray) of myrB. HDV infection was monitored by detecting intracellular HDV RNA (left graph) and HDAg (right pictures) by real time RT-PCR and immunofluorescence, respectively. In the lower pictures, red and blue signals indicate HDAg and nucleus, respectively. Scale bar, 100 µm.

### Sequencing of NTCP in cat liver and the database of whole genome sequence in lion

To validate the reference sequence of cat NTCP registered in the NCBI RefSeq database (*Felis catus*, XM_003987782.5), we analyzed whole-genome assemblies from four cat individuals (GCA_000003115.1, GCA_000181335.6, GCA_013340865.2, and GCA_016509815.2) as shown in Methods, and found these seqnences have no or one base variation but are completely identical at amino acid level to the reference cat NTCP sequence (S3A and B Fig). We further recovered total RNA from the liver of a domestic cat in Japan to determine the sequence of its cDNA, and confirmed the resulting cDNA sequence was fully identical to the reference cat NTCP sequence (S3A and B Fig).

Using the same approach, we analyzed whole-genome assemblies from four lion individuals (GCA_008795835.1, GCA_012313985.1, GCA_046254995.1, and GCA_046255005.1) to infer NTCP sequences, and found no to two base variations but are completely identical at amino acid level to the reference lion NTCP sequence (S3C and D Fig). As we cannot access to sample the liver derived from a lion, we validated the amino acid residues at aa 191 and 268 (as key residues identified, as shown below) in the lion NTCP reference sequence by analyzing whole-genome sequencing reads derived from Sequence Read Archive (SRA) from 10 lion individuals as well as 1 leopard as a reference (10 Lions [Panthera leo] and 1 Leopard [Panthera pardus])[40], as shown in Methods (S3E Fig). The inferred amino acid resides at aa 191 and 268 in NTCP were consistent with those in the reference lion NTCP (R191 and A268) in all individuals with available sequence information, although aa 191 could not be determined in one individual and aa 268 in another. In contrast, the corresponding residues in the leopard matched the reference leopard NTCP sequence (G191 and T268) (S3E Fig).

Thus, we confirmed the reference amino acid sequences of the cat and lion NTCPs employed in this study are not restricted to a single individual, but are representatives of multiple individuals within each lineage.

### Amino acid substitution at aa 191 and 268 in cat NTCP impairs the capacity for supporting preS1 binding and viral infection

We analyzed the amino acids responsible for the distinct viral receptor capacity among feline familly-derived NTCPs. Seven amino acid sequences (aa 121, 191, 195, 216, 245, 268, and 283) in lion NTCP are unique compared with NTCPs derived from cat, tiger, and leopard (Fig. 2A). To identify amino acid essential for viral receptor function, we substituted each amino acid in cat NTCP to its lion counterpart residue, and overexpressed them in HepG2 cells to evaluate preS1 binding. As shown in Fig. 4A, substitution at aa 191 in cat NTCP significantly reduced preS1 binding levels to 29–44% of the wild type cat NTCP, and that at aa 268 more drastically reduced preS1 binding to less than 10% (Fig. 4A, lanes 1, 3, and 7). Combined substitution at aa 191 and 268 completely abrogated the preS1 binding activity of cat NTCP at any preS1-treated concentrations (Fig. 4A, lanes 9). Similar trend was observed in HBV infection assay using HepG2 cells overexpressing the cat NTCP or its mutants (Fig. 4B). Expression levels of these NTCP variants on cell surface were not much variable (Fig. 4C), suggesting that the effects of amino acid substitution are not due to the altered cell surface expression of NTCP. These data suggest that G191 and T268 in cat NTCP are critical for its function for supporting preS1 binding and viral infection.

**Fig. 4.**
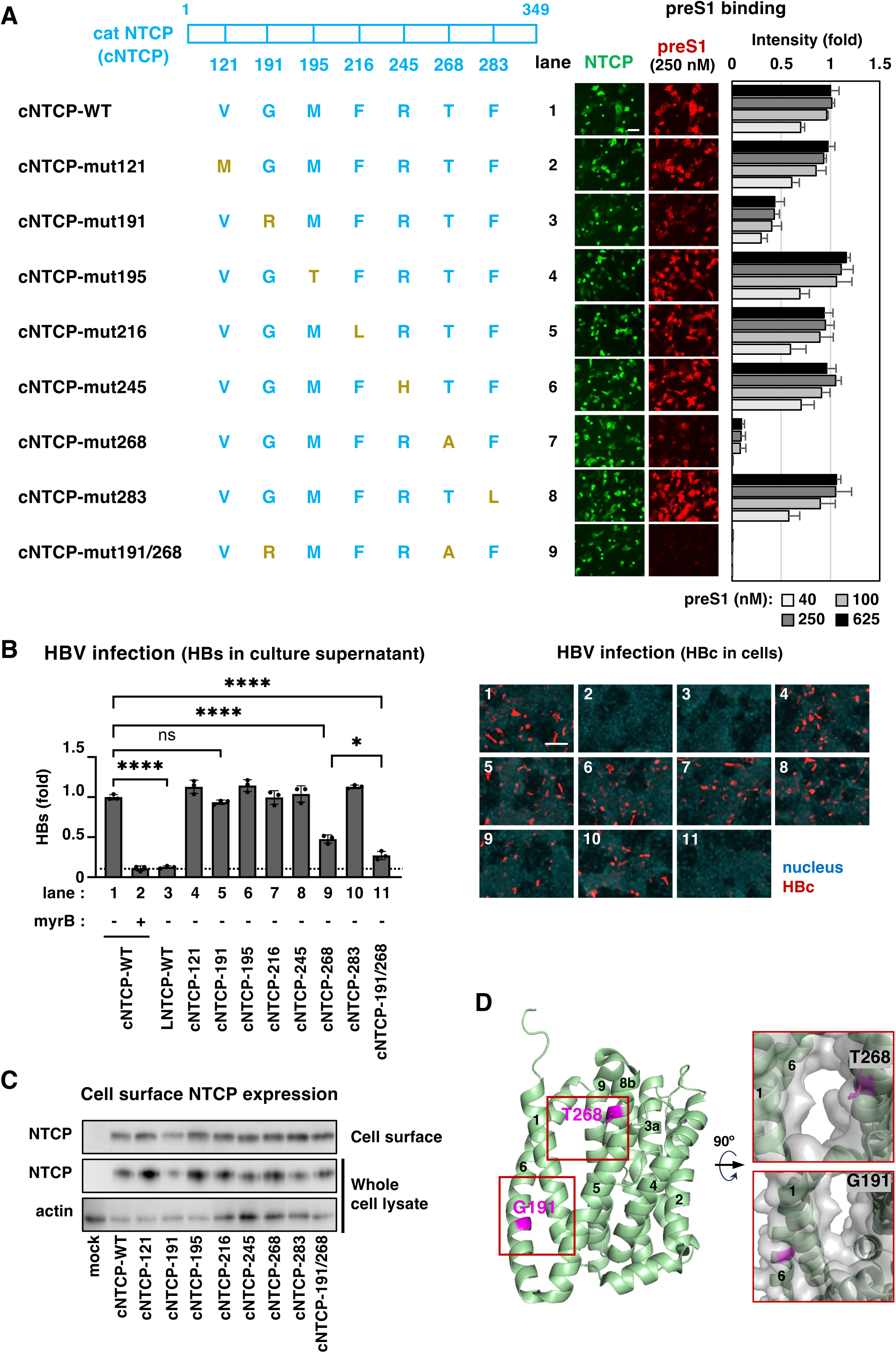
Identification of amino acids in cat NTCP important for supporting preS1 binding and HBV infection. **(A)** Left, schematic diagram of the wild type and the amino acid-substituted variants of cat NTCP (cNTCP) are shown. The amino acids derived from cat and lion are shown by blue and yellow, respectively. Right, Capacity of preS1 binding assessed by preS1 binding assay are shown. The pictures indicate the cell-bound preS1 (red) as well as NTCP (green) in HepG2 cells overexpressing the indicated NTCPs upon treatment with 250 nM preS1-TAMRA in the preS1 binding assay. The right graph shows the preS1 binding activity calculated by dividing preS1 (red) intensity by NTCP (green) area, to each NTCP cells for varying amount of preS1-TAMRA (40, 100, 250, and 625 nM). **(B)** HBV infection assay were performed using HepG2 cells overexpressing each cNTCP variant or the wild type either in the presence (+) or absence (-) of myrB, as shown in Fig. 2B. Extracellular HBs (left) and intracellular HBc (right pictures) are shown. Red and blue signals in the right pictures indicate HBc and nucleus, respectively. Scale bar, 100 µm. **(C)** Expression levels of each cNTCP on cell surface and in total cell fraction were detected by immunoblot, as shown in Fig. 2D. **(D)** Left, positions at 191 and 268 are indicated (magenta) based on the cryo-electron microscopy structure of human NTCP with ribbon diagram (lime). Transmembrane 1–9 are shown. Right, high resolution images around G191 and T268 are shown.

The aa 191 and 268 are located at TM6 and TM8b, respectively, based on the cryoEM structure of human NTCP, which we have recently reported[24] (Fig. 4D). The aa 268 faces on the bile acid tunnel, closely contacting to preS1 in the cryoEM structure of the preS1/human NTCP complex, whereas aa 191 resides close to TM1 to form the rigid panel domain, but not exposed on the bile acid tunnel surface[26,27].

### Combined substitution at aa 191 and 268 in lion NTCP acquires the viral receptor function

Conversely, we substituted the seven amino acids of the lion NTCP-backbone with the cat-type residues to examine the capacity of preS1 binding and HBV infection. As shown in Fig. 5A, simultaneous substitution of all the seven amino acids in lion NTCP to the cat-type counterparts (lNTCP-7mut) acquired the preS1 binding equivalent to the levels for the wild type cat NTCP (Fig. 5A, lane 10). However, any single substitution in lion NTCP among the seven amino acids did not show any visible preS1 binding (Fig. 5A, lanes 2–8). In contrast, the double substitution of lion NTCP at aa 191 and 268 (lNTCP-191/268) restored the marked preS1 binding to almost the equivalent level to the wild type cat NTCP (Fig. 5A, lane 9), suggesting that these two amino acids, in combination, are required and sufficient for acquiring the preS1 binding capacity. Also in the HBV infection assay using HepG2 cells overexpressing each lion NTCP variant, the double substitution at aa 191 and 268, as well as the seven-point substitution, acquired the HBV infection (Fig. 5B). This infection assay showed a slight infection in the single mutant at aa 268, which however was much lower than that with cat NTCP (Fig. 5B, lane 9). Similarly to Fig. 4C, we also detected the cell surface expression of these NTCPs, showing no remarkable alteration in the expression level and cell surface localization by amino acid substitutions at either aa 191 or 268 (Fig. 5C). These results suggest that double substitutuion at aa 191 and 268 determines the functionality of feline-derived NTCPs as a viral receptor.

**Fig. 5.**
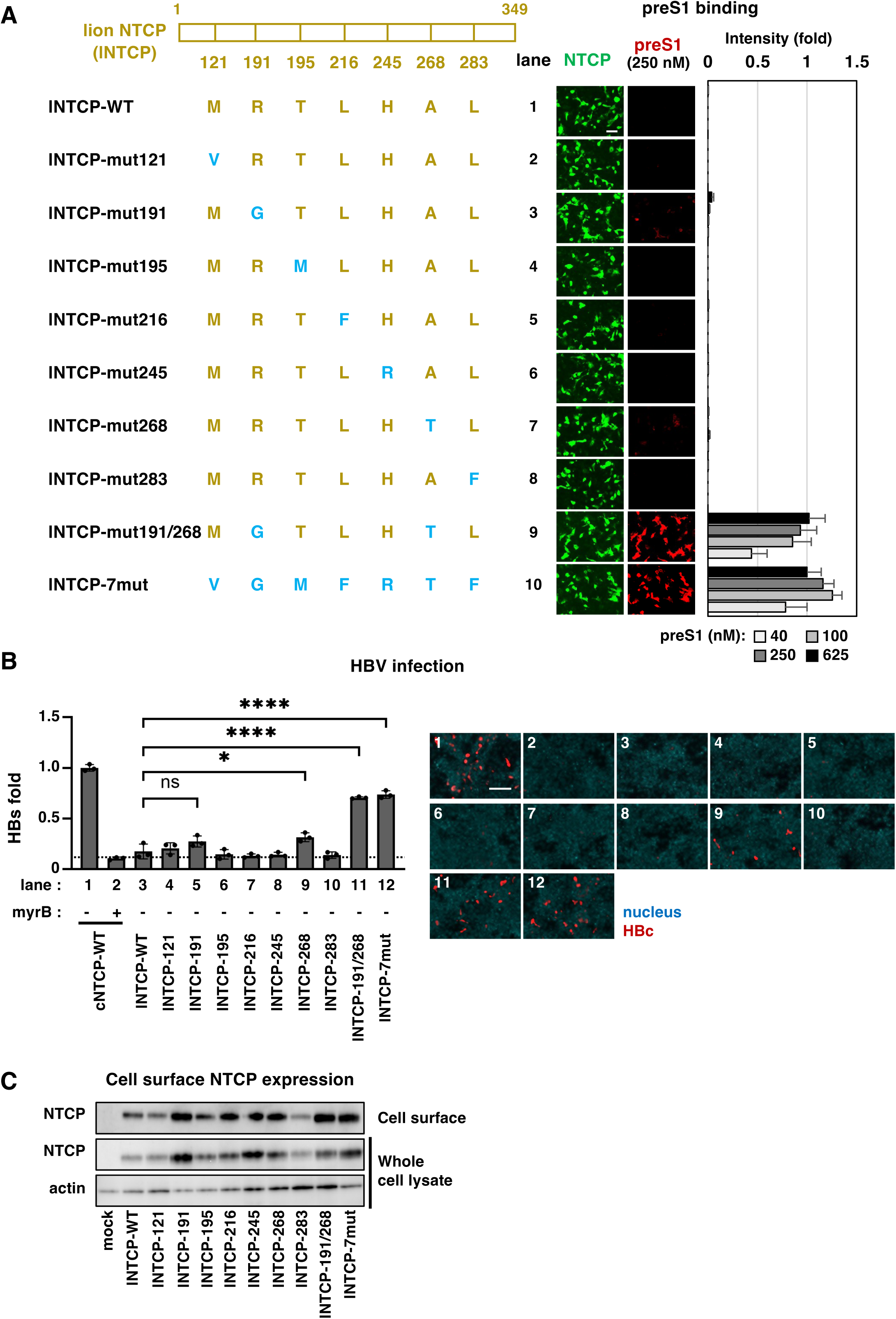
Gain of viral receptor function of lion NTCP by amino acid substitution to the cat counterparts. **(A)** Left, schematic diagram of the wild type (yellow) and the variants of lion NTCP (lNTCP), of which amino acids were substituted to the cat counterparts (blue). Right, Immunofluorescence images and the graphs for preS1 binding activity are indicated as shown in Fig. 4A. **(B)** Capacity of the indicated NTCPs for supporting HBV infection was examined by HBV infection assay as shown in Fig. 4B. Left graph and right pictures indicate extracellular HBs and intracellular HBc, respectively. Red and blue signals in the right pictures indicate HBc and nucleus, respectively. Scale bar, 100 µm. **(C)** Expression levels of the indicated NTCPs on cell surface and in total cell fraction were assessed by immunoblot as shown in Fig. 2D.

### Molecular dynamics simulations suggest the distinct dynamic structure of lion NTCP compared with that of human or cat NTCP

We tried to solve the cryoEM structure of cat and lion NTCPs, however, it was unsuccessful because of the technical difficulty. Alternatively, to get an insight into the mechanism for the distinct viral receptor function among lion- and cat-derived NTCP, we performed molecular dynamics (MD) simulations.

We prepared the structural models of cat and lion NTCPs prepared by homology modeling based on the cryoEM structure of human NTCP in complex with preS1 (PDB: 8HRY)[26], from which preS1 was removed prior to MD simulations, yielding apo-like starting structures. Differences in conformations among these NTCPs were analyzed by principal component analysis of MD trajectories (Figs 6A, and S4 Fig). Structural distributions of human and cat NTCPs remained around the experimental cryoEM structures of human NTCP, i.e. apo human NTCP at outward-facing state (PDB: 7FCI, diamond), human NTCP complexed with HBV preS1 (PDB: 8HRY, triangle), and that in complex with bile acids (PDB: 7ZYI, circle) (Fig. 6A, upper and middle), whereas lion NTCP sampled structural distributions far from these cryoEM structures, with three divided basins (Fig. 6A, lower). Despite starting from nearly identical structures in the MD simulations, lion NTCP particularly showed distinct structural distibutions: Lion NTCP, without structural disruption, maintained the overall human NTCP-like structure, but showed variations in the relative positioning and axis straightness of the TM helices. Especially, TM6 was nearly linear with 170–180 degrees of the bending angle at residue 191 in human and cat NTCP, whereas TM6 in lion NTCP bent by 20–30 degrees to the inner bile acid tunnel side, compared to cat NTCP (Fig. 6B, yellow v.s. cyan or green). Thereby, the neighboring TM1 was tilted toward TM5 in lion NTCP, with a reduced distance between TM1 and TM5 (as shown by the distance between Cα at residue 31 and Cα at residue 162) by approximately 2 Å (Fig. 6C, lower left). These TM repositionings altered the width of the preS1 binding pocket within the bile acid tunnel: The crevice between TM5 and TM8b (as shown by the distance between Cα at residue 162 and Cα at residue 264), which is the most critical regions determining the preS1 binding capability of NTCP[41], was 14–20 Å in lion NTCP, substantially larger than 8–12 Å observed in human and cat NTCPs (Fig. 6C, lower right). These structural features can explain the lack of the preS1 binding capability of lion NTCP. To further examine whether the observed structural differences in lion NTCP are related to the loss of functionality as an HBV receptor, we performed MD simulations using the lion NTCP carrying substitutions at residues 191 and 268 to the cat counterparts [lion NTCP(R191G/A268T)], which rescued the viral receptor function. As shown in Fig. 6D, substitutions at residues 191 and 268 to the cat counterparts in lion NTCP approached to the cat NTCP-like distribution that was more frequently close to the experimental cryoEM structures of human NTCP (Fig. 6D, lower). Linearity of TM6 in lion NTCP became closer to that in cat NTCPs by the cat-mimic substitutions at residues 191 and 268 (Fig. 6E, red v.s.cyan). Consistently, the bile acid tunnel region, one of the critical regions determining the preS1 binding capability (crevice between TM5 and TM8b), was narrowed and almost identical to that of cat NTCP by the 191/268 substitution to the cat counterparts in lion NTCP (Fig. 6F, red v.s. cyan).

**Fig. 6.**
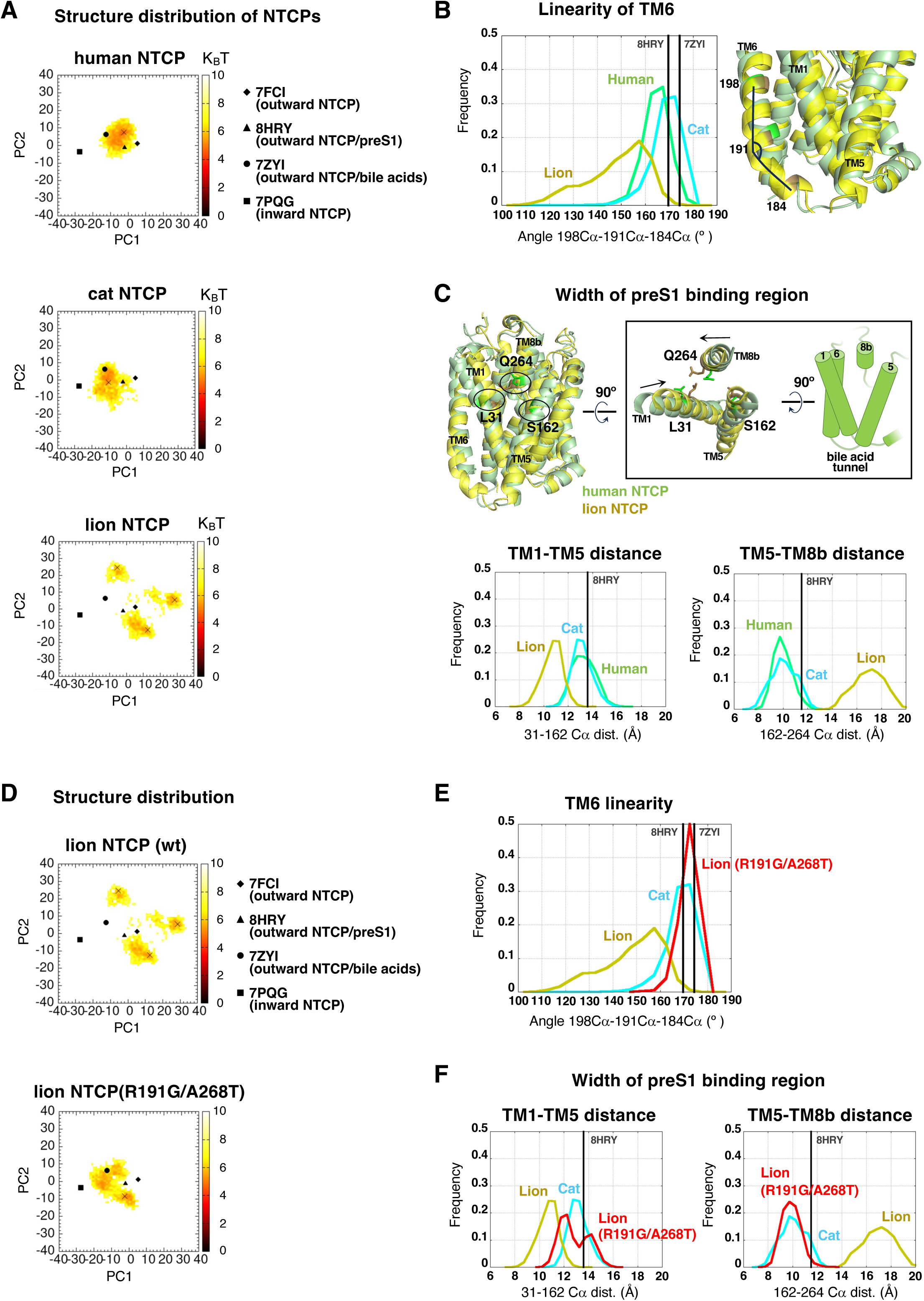
Molecular dynamics simulations indicating the different structural features between cat and lion NTCPs. **(A)** Structural distributions on principal component (PC) surface for apo state of human NTCP (upper), cat NTCP (middle), and lion NTCP (lower). Frequencies are shown by color (Brown to white, high to low). The points for the most frequent distribution are shown by x. Filled diamond, triangle, circle, and square indicate the cryoEM structure of the outward facing form of apo human NTCP (PDB: 7FCI), the outward facing human NTCP in complex with preS1 (PDB: 8HRY), the outward facing human NTCP complexed with bile acids (PDB: 7ZYI), and the inward facing apo human NTCP (7PQG), respectively. **(B)** Left, Frequency of TM6 bending angle during the three independent MD simulations for human (green), cat (cyan), and lion (yellow) NTCP systems. The angles for cryoEM structure of NTCP in complex with preS1 (PDB: 8HRY) and that in complex with bile acids (PDB: 7ZYI) are shown by the black vertical lines as references. Right, The definition of the TM6 bending angle is shown by black line at 191C_α_. **(C)** Altered structure of the preS1 binding region in the bile acid tunnel. Upper, Overlay structure of human and lion NTCP is shown by green and yellow, respectively. The right inset indicates the top view of TM1, 5, and 8b (left) and the side view of TM1, 5, 6, and 8b (right). Lower, Frequency of the distance between C_α_ atom at position 31 and that at position 162 (left, for TM1-TM5 distance), and that between C_α_ atom at position 162 and that at position 264 (right, for TM5-TM8b distance). **(D)** Structural distributions on PC surface for the wild type lion NTCP (upper) and lion NTCP(R191G/A268T) (lower) systems, as shown in (a). **(E)** Frequency of the TM6 bending angle for cat NTCP (cyan), lion NTCP (yellow), and lion NTCP(R191G/A268T) (red), as shown in (b). **(F)** Frequency of the width of the preS1 binding region for cat NTCP (cyan), lion NTCP (yellow), and lion NTCP(R191G/A268T) (red), defined by distance between 31–162 residues (left, for TM1-TM5 distance) and between 264–162 residues (right, for TM5-TM8b distance), as shown in (c).

The above calculations have a limitation in that they are simulations using modeled structures based on the cryoEM human NTCP structure, rather than the experimental structures of cat and lion NTCPs. However, our computational data are readily in a good agreement with that NTCP derived from cat as well as human maintains a structure favorable to preS1 binding, whereas lion NTCP has unique structural behavior with a wider TM5-TM8b crevice (8–12 Å in human and cat NTCPs v.s. 14–20 Å in lion NTCP) that is likely unfavorable to stable preS1 binding.

### Diverse functionality of feline-derived NTCPs determines the susceptibility to domestic cat HBV

Hepadnavirus originally infected in domestic cat (domestic cat HBV, DCHBV) has been reported[7]. To analyze the feline-derived NTCP functionality as a DCHBV receptor, we examined the binding of a fluorescence-labelled preS1 DCHBV 2–48 aa peptide (Fig. 7A, left). In this binding assay, HepG2 cells overexpressing NTCPs derived from cat, tiger, and leopard showed the binding of DCHBV preS1 in a concentration-dependent manner, and these signals disappeared in the presence of excess amount of either non-fluorescence labelled DCHBV preS1 or myrB (HBV-preS1) (Fig. 7B, lanes 1–3). In contrast, cells expressing lion NTCP did not support binding at any treated concentrations of fluorescence-labelled DCHBV preS1 (Fig. 7B, lane 4). Furthermore, substitution at both aa 191 and 268 in cat NTCP with the lion residues completely abolished binding activity of DCHBV-derived preS1 (Fig. 7B, lane 5), and inversely, the lion NTCP variant with dual aa 191 and 268 substitution or the seven amino acid substitution acquired DCHBV preS1 binding (Fig. 7B, lane 6 and 7), consistent with the results in Fig. 4 and 5. These data suggest that DCHBV preS1 binds to NTCP in a manner similar to that of HBV preS1. We then confirmed the susceptibility of these NTCP-expressing cells to HDV-based pseudovirus carrying the DCHBV preS1^2–48^ region, to evaluate DCHBV preS1-mediated entry (Fig. 7A, right). As shown in Fig. 7C, HepG2 cells overexpressing NTCPs derived from cat, tiger, leopard, and lion NTCPs carrying either the aa 191/268 double replacement or the seven mutations showed intracellular HDV RNA detection, which were reduced by treatment with myrB to the background level (Fig. 7C, lanes 1, 2, 3, 6, and 7). Consistent with the results above, the cells overexpressing lion NTCP and cat NTCP with the lion-type 191/268 substitution were resistant to HDV infection (Fig. 7C, lanes 4 and 5). These results suggests that the sequence at aa 191 and 268 of feline NTCPs are determinant of DCHBV susceptibility.

**Fig. 7.**
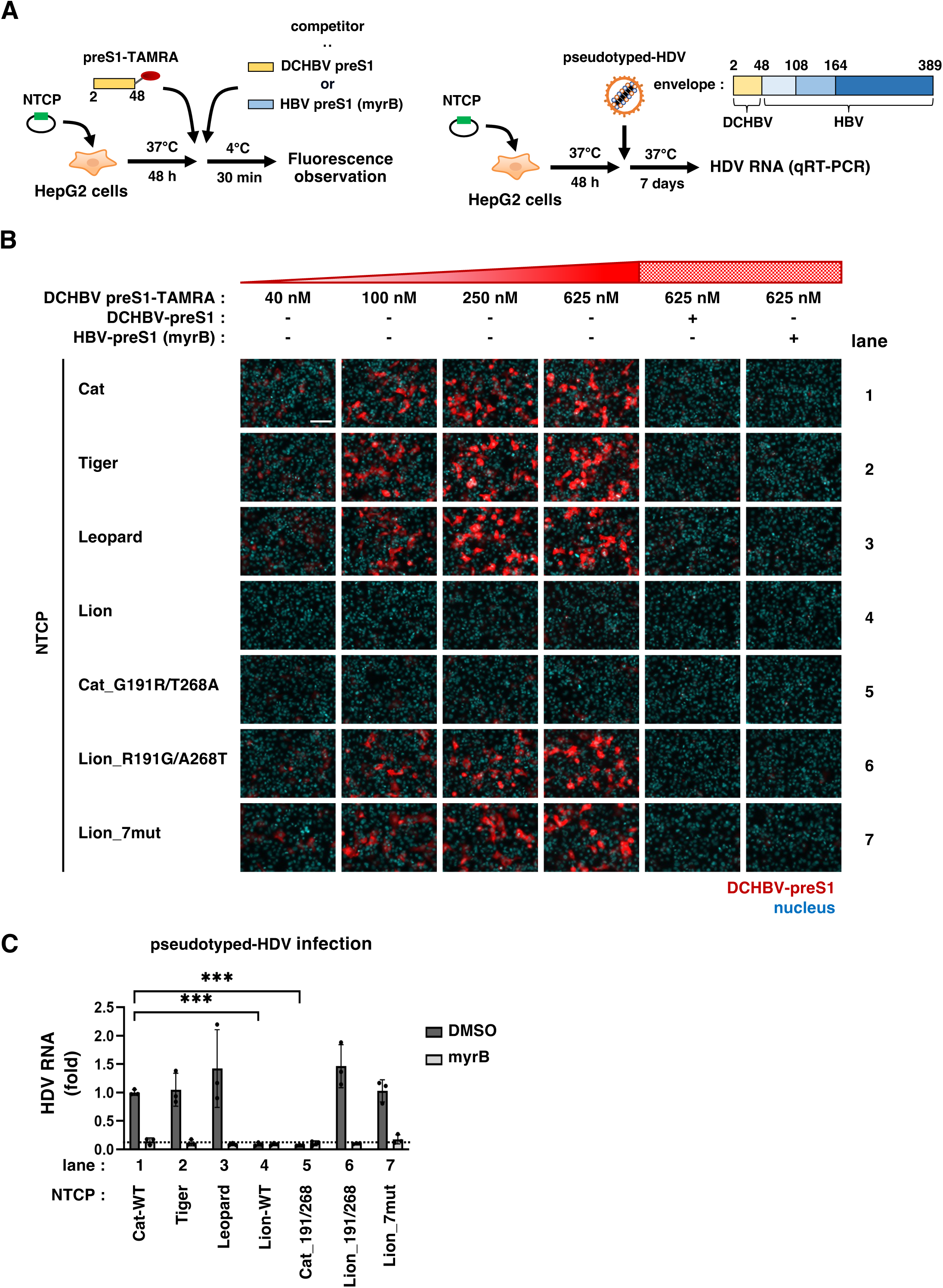
Capacity of feline family-derived NTCPs as a receptor for domestic cat HBV (DCHBV). **(A)** Scheme for the binding assay of DCHBV-derived preS1(2–48) conjugated with TAMRA (left) and infection assay of pseudotyped HDV carrying the DCHBV-derived preS1(2–48) (right), essentially perfomed as shown in Fig. 1B and C. **(B)** HepG2 cells transfected with the expression plasmid for the indicated NTCPs were subjected to the preS1 binding assay using the TAMRA-labelled DCHBV preS1 peptide (DCHBV preS1-TAMRA) at 40, 100, 250 or 625 nM, as shown in a, left. Binding assay with 625 nM DCHBV preS1-TAMRA in the presence of non-TAMRA labelled DCHBV preS1 or HBV preS1 was also shown for competition (The right two panels). The immunofluorescence detection images are shown. Red, DCHBV preS1; blue, nucleus. Scale bars, 100 µm. **(C)** HepG2 cells overexpressing the indicated NTCPs were subjected to the pseudotyped-HDV infection assay, as shown in a, right. The assay was performed both in the presence (light gray) or absence (dark gray) of myrB and evaluated by detecting intracellular HDV RNA by real time RT-PCR.

## Discussion

In this study, we showed that despite being as much as 97% identical in amino acid sequence, NTCPs derived from feline species are non functionally equivalent in HBV/HDV receptor capacity. Accumulating evidences so far show that NTCP is a major determinant for host cell susceptibility to HBV and HDV infection, mainly through distinct preS1 binding capability. It was recently revealed that the bile acid tunnel and the outer-surface hollow in NTCP play a central role in mediating preS1 binding through a multistep interaction mechanism[23–29]. These two sites are also involved in viral restriction in distinct animal species: The 84–87 aa (SDM-1) in the outer-surface hollow and 157–165 aa (SDM-2) in the bile acid tunnel (and their surrounding regions) display high sequence diversity among species and determine the host susceptibility to infection[5,6,18,32–34] (Fig. 8-i). In SDM-2, substitution of aa 158 (glycine in human NTCP) to longer-side chain amino acid (macaque and pig NTCP) forms a steric hindrance at the entrance of the bile acid tunnel that prevents preS1’s insertion into the tunnel[29] (Fig. 8-ii, macaque & pig). In SDM-1, the 84–87 aa of mouse NTCP (HLTS in mouse v.s. RLKN in human) forms an outer-surface hollow having lower preS1 binding compatibility, and hence resistance to HDV infection[19,32] (Fig. 8-iii, mouse). A substitution at the amino acid surrounding position 82 abolished the virus receptor function of cow and goat NTCP (F82 in cow and goat v.s. V82 in horse and human)[42] (Fig. 8-iii, cow). Importantly, these two motifs are involved specifically in preS1 binding but not in bile acid transport, therefore an evolutationary hot spots of mutation that drives high diversity of viral susceptibility among species but preserves the bile acid transporter function. The particular function of lion NTCP reported here is in clear contrast to these reports of previous studies: Substitution of the amino acid residues at aa 191 and 268 suggests reconstructig the overall structure of the NTCP bile acid tunnel, with marked reduction in the bile acid transporter function in lion NTCP (Fig. 8-iv, feline family). This indicates that evolutionary genetic change allows functional impairment of bile acid transporter function of NTCP. How does the lion maintain bile acid homeostasis under the impairment of NTCP function? As another bile acid transporter family, organic anion transporters (OATP)s are expressed on the basolateral membrane of hepatocytes[43], these transporters may complement, at least partly, the bile acid transport function.

**Fig. 8.**
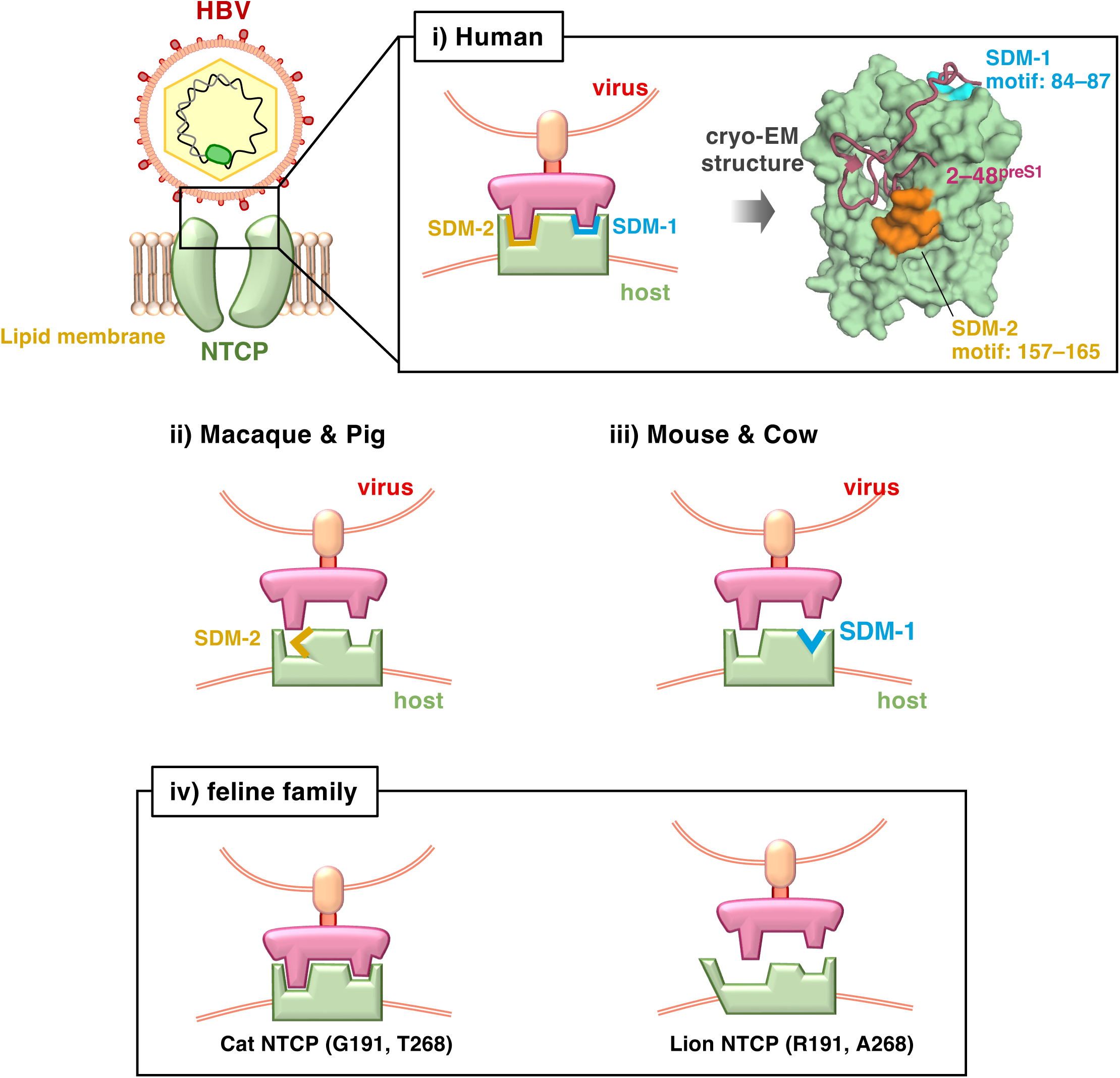
Determinants in NTCP for species susceptibility to hepadnavirus infection. i) In human NTCP, SDM-1 (motif: 84–87 aa, orange) and -2 (motif: 157–165 aa, cyan) constitute the interface to preS1(2–48) (red) at the extracellular surface and bile acid tunnel, respectively. ii) In macaque and pig, steric hindrance formed by amino acid substitutions in SDM-2 is the major determinant for non-susceptibility to HBV/HDV infection. iii) In mouse and cow NTCP, amino acid substitutions in SDM-2 (and the peripheral region) are responsible for their deficientcy for viral receptor function. iv) This study suggests that the overall structural diversity of the bile acid tunnel in feline species (determined by positions 191 and 268), independently of SDM-1 and -2, determines the NTCP functionality to support host susceptibility to hepadnavirus infection.

Based on the cryoEM structure of the human NTCP in complex with preS1^2–48^ [26,27], T268 in NTCP is located on TM8b at the entrance of the bile acid tunnel and closely contacts with the multiple amino acids of preS1^2–48^, especially N4^preS1^, S6^preS1^, P15^preS1^, and W32^preS1^ [26]. Its substitution with the lion-type alanine (having shorter side chain), would impair the interaction energy with these preS1 amino acids. In contrast, aa 191 is located within TM6 and does not directly contact to preS1, but it is adjacent to TM1 (F37^NTCP^ and S41^NTCP^) that resides in parallel with TM6 to constitute the panel domain of NTCP. Substitution of small glycine (cat- and human-type) to the long side-chain and positive-charged arginine (lion-type) at aa 191 was suggested to change the linearity of TM6 by our MD simulations, in which TM6 in cat NTCP is almost linear (tilt angle=170–180°), whereas that in lion NTCP is bent with sharp tilt (tilt angle=150°–160°) to the inner side (Fig. 6B) that would induce local steric hindrance to the neighbor TM1 and reposition the structure of the bile acid tunnel. By this repositioning, the crevice between TM5 and TM8b, as the bile acid tunnel region predominantly determining the preS1 binding capability[41], was simulated to be far wider in lion NTCP, with nearly double the size of that of human or cat NTCPs (14–20 Å in lion NTCP v.s. 8–12 Å in human and cat NTCPs) (Fig. 6C), likely disabling high affinity preS1 insertion. Future cryoEM analysis is expected to demonstrate experimentally the difference in the bile acid tunnel structure of cat and lion NTCPs, but our simulations already well explain the basis how the functionality of feline-derived NTCPs is diverse, depending on the structure of the bile acid tunnel.

DCHBV was recently identified as an HBV-related orthohepadnavirus originally infected in domestic cats with chronic hepatitis, with a prevalence of 0.8–25% in the domestic cat populations depending on countries[7]. We have reported that DCHBV infects host hepatocytes in an NTCP-dependent manner[37]. However, the significance of cat NTCP in determining the host species of DCHBV as well as its host range itself remains undefined. The present study suggests that NTCPs derived from cat, tiger, and leopard, but not lion, function as receptors of DCHBV in a aa 191 and 268-dependent manner, similarly to HBV/HDV. This is the first report indicating the molecular basis whereby DCHBV recognizes and selects host species. We suggest that only a limited subset within the Feline family can be infected by DCHBV, consistent with the general hypothesis that hepadnaviruses have narrow host range. It should be also noted that the host susceptibility to hepadnaviruses is governed not only by virus binding but also by post-binding process including internalization and membrane fusion.

The present study showed that primary cat hepatocytes could be infected with HDV. Limitation of our study is the lack of examination for HBV infection in primary cat hepatocytes due to technical difficulties, which should be optimized in the future. Recently, however, another group has reported the successful infection of primary cat hepatocytes with HBV and HDV[39], consistent with our conclusion that cat NTCP is functional as an HBV and HDV receptor. In their publication, it remains unclear how the HBV/HDV susceptibility is determined among primary hepatocytes derived from many species. We analyzed here that the infection susceptibility among feline species is determined by NTCP depending on the structure of the bile acid tunnel.

In conclusion, we identified a mechanism for regulating the viral receptor functionality of NTCP and determining the susceptibility to HBV/HDV infection that involves coordinately the two amino acids at distinct transmembrane regions. Among feline species, lion NTCP is unique in escaping from viral infection but unexpectedly losing the original function for bile acid uptake in nature. Our evidences provide an additional aspect for mechanism determining the host range of HBV and related hepadnaviruses.

## Methods

### Reagents

Biotinylation reagent (EZ-Link^TM^ Sulfo-NHS-LC-Biotin) and Streptavidin beads (Streptavidin Agarose Resin) were purchased from Thermo Fisher SCIENTIFIC. Myristoylated preS1 peptides consisting of 2–48 aa region derived from HBV and DCHBV labelled with TAMRA (preS1-TAMRA)[37] were synthesized by Scrum.

### Cell Culture

HepG2 cells were cultured in DMEM/F12+Glutamax supplemented with 10 mM HEPES, 100 units/ml penicillin, 100 µg/ml streptomycin, 10 % fetal bovine serum (FBS; NICHIREI), and 5 µg/ml insulin[44]. Huh-7 cells were maintained in D-MEM (High Glucose) with L-Glutamine and Phenol Red containing 10 % FBS, 10 mM HEPES, 100 units/ml penicillin, 100 µg/ml streptomycin, 0.1 mM nonessential amino acid, and 1 mM sodium pyruvate. Primary culture of rat (BIOIVT) and cat (Visikol) hepatocytes were cultured according to the manufacturing instruction. All cells were cultured at 37°C in 5% CO_2_.

### Construction of plasmids and DNA transfection

Expression plasmids of mammalian NTCPs, which were inserted into the KpnI or BamHI and XbaI sites of the pEF4-Myc/His, were generated by Genscript. All mutant constructs were prepared by oligonucleotide-directed mutagenesis essentially as described[29]. The pT7HB2.7-DCHpreS1 plasmid, encoding the HBV LHBs, of which 2-48 aa region of preS1 was replaced by that of DCHBV, was generated by oligonucleotide-directed mutagenesis using pT7HB2.7, an expression plasmid for HBV LHBs, as a template[45].

HepG2 cells were transfected with each plasmid using Viafect (Promega) according to the manufacture’s protocol and were subjected to the assays at 48 h posttransfection. For producing HDV or HDV-based pseudovirus, Huh-7 cells were transfected with pSVLD3 (kindly provided by Dr. John Taylor at Fox Chase Cancer Center) and pT7HB2.7 or pT7HB2.7-DCHBVpreS1 using lipofectamine 3000 (Thermo Fisher SCIENTIFIC), according to the manufacture’s protocol.

### PreS1 binding assay

NTCP activity to support preS1 binding was evaluated by incubating with the indicated concentrations of the N-terminally myristoylated preS1 peptide consisting of 2–48 aa region conjugated with TAMRA fluorescence (preS1-TAMRA) at 4°C for 30 min. After washing, the cells were fixed with 4% paraformaldehyde to simultaneously detect NTCP and the nucleus as described in the following method for immunofluorescence analysis[6]. Fluorescence signals for preS1, NTCP, and the nucleus were visualized with a fluorescence microscopy, BZ-X810 (Keyence). The signals were quantified with BZ-H4A and BZ-H4C (Keyence) to measure the fluorescence intensity of cell-bound TAMRA as well as the area of NTCP signals and calculated as preS1 susceptibility of NTCP to show as a graph (Fig. 1, 4A, and 5A). The background level for the preS1-TAMRA signal attached to cells was shown under treatment with excess amount of myrcludex-B, a preS1 attachment inhibitor.

### Immunofluorescence analysis

The fixed cells were permeabilized with 0.01% digitonin or 0.3% Triton X-100 for 10 min, followed by incubation with either anti-Myc (Santa Cruz), anti-pan-cadherin (Abcam), anti-HBV core antigen (synthesized from Biogate), or anti-HD antigen (synthesized from Scrum) as primary antibodies for 16 h. After washing, the cells were incubated with Alexa488- and Alexa555-conjugated secondary antibodies together with 0.02% DAPI for 1 h. Fluorescence image were obtained using BZ-X810. In S2A Fig, subcellular localization of NTCP was analyzed with confocal microscopy, LSM 900 with Airyscan 2 (ZEISS).

### Detection of cell surface proteins

The cells were treated with 0.5 mg/mL of biotinylating reagent (Thermo Fisher SCIENTIFIC) at 4°C for 30 min and then washed out excess reagent to biotinylate cell surface proteins. The cells were lysed with the buffer [50 mM Tris-HCl (pH 7.6), 150 mM NaCl, 5 mM EDTA, 1% NP-40, and complete protease inhibitor] and pulled down with streptavidin beads (Thermo Fisher SCIENTIFIC). The precipitates were loaded by SDS-PAGE and detected by immunoblot analysis essentially as described[6] with anti-Myc (Santa Cruz) and anti-actin (Sigma) antibodies as primary antibodies. For NTCP detection, the precipitate samples were treated with 250 U Peptide-N-Glycosidase F (PNGase F) to digest N-linked oligosaccharides from glycoproteins before SDS-PAGE[29]. As a control, total cell lysate fraction without pulldown with streptavidin beads was also detected with anti-Myc and anti-actin antibodies to visualize the intracellular expression levels.

### Transporter assay

NTCP-dependent bile acid uptake activity was measured essentially as described[6]. The cells were treated with [^3^H]-taurocholic acid as a substrate in either sodium-containing or -free buffer at 37°C for 10 min. After washing out free [^3^H]-taurocholic acid, the cells were lysed with 0.05% SDS and the intracellular radioactivity was measured using liquid scintillation counter (PerkinElmer). NTCP-mediated transporter activity was evaluated by uptake of radioactivity upon sodium-containing buffer.

### Infection assay

HBV used in the inoculum for the HBV infection assay was derived from the culture supernatant of a tet-off cell line for HBV expression, Hep38.7-Tet cells, under depletion of tetracycline, as described[46]. The cells were inoculated with HBV in the presence of 4% PEG8000 for 16 h, which were then washed and incubated for an additional 12 days to detect HBs antigen in the culture supernatant and HBc antigen in the cells by ELISA and immunofluorescence analysis, respectively, as described[6].

Inoculum of HDV or HDV-based pseudovirus were recovered from the culture supernatant of Huh-7 cells transfected with pSVLD3 together with either pT7HB2.7 or pT7HB2.7-DCHBVpreS1, as described[47]. The cells were inoculated with these viruses for 16 h and then washed, incubated for 6 days, and harvested to detect HDV RNA and HD antigen in the cells by real time reverse transcription (RT)-polymerase chain reaction (PCR) and immunofluorescence analysis, respectively, as described[48].

### ELISA

The recovered cell culture supernatants were applied to anti-HBs antibody-conjugated plates, followed by washing and incubating with horseradish peroxidase-conjugated anti-HBs antibody to detect with TMBZ reagent, as described previously[6].

### Real time RT-PCR

Intracellular RNA was extracted using RNeasy mini kit (QIAGEN) according to the manufacture’s protocol. HDV RNA were measured by real time RT-PCR analysis with THUNDERBIRD Probe One-step qRT-PCR Kit (TOYOBO) using 5’-GGACCCCTTCAGCGAACA-3’, and 5’-CCTAGCATCTCCTCCTATCGCTAT-3’ for the primer sets and 5’-FAM-AGGCGCTTCGAGCGGTAGGAGTAAGA-TAMRA-3’ for the probe, as described[48].

### Phylogenetic analysis

Phylogenetic analysis was performed using amino acid sequence of NTCP from 23 species (Supplementary Table 1) in the public database. Multiple sequence alignment was conducted using MAFFT with default parameters[49,50]. The phylogenetic tree was constructed using the Neighbor-Joining method implemented in MEGA 11[51]. Bootstrap analysis was performed with 1,000 replicates to assess the statistical support for the tree.

### Validation of NTCP sequences

Cat liver collection was performed as follows. The uterus was obtained from a cat that underwent ovariohysterectomy for medical reasons at a university-affiliated veterinary hospital in Japan and was kept on ice until use. The fetuses were already dead at the time the uterus was collected. Therefore, no animals were euthanized for the purposes of this study, and no euthanasia method or anesthetic agent for fetal sacrifice was applicable. All procedures were approved by the Animal Care and Use Committee of the University of Miyazaki (Approval No. 2023-012) and were conducted in accordance with the University of Miyazaki experimentation regulations and relevant institutional guidelines. Liver tissue was disrupted using a Biomasher (Nippi) and were lysed in Buffer RLT supplemented with 2-mercaptoethanol (Bio-Rad). RNA was subsequently purified according to the RNeasy Mini Kit protocol (Qiagen). The coding sequence of NTCP was amplified using PrimeScript II High Fidelity One Step RT-PCR Kit (TaKaRa) according to the manufacture’s protocol, using forward (5’-ACCCAGTTGCATAATCCTTCTATTT-3’) and reverse (5’-TTGTCAGTTCTGATAGTTTCTGCTG-3’) for the primer sets. The PCR fragments were used for a direct sequencing.

To validate the NTCP sequence derived from domestic cat registered in the NCBI RefSeq database (*Felis catus,* XM_003987782.5), we analyzed whole-genome assemblies from four individuals (GCA_000003115.1, GCA_000181335.6, GCA_013340865.2, and GCA_016509815.2). Publicly available data were downloaded in FASTA (.fna) and formatted into searchable BLAST databases using the *makeblastdb* from the BLAST v.2.11.0. To identify homologs of the cat NTCP sequence, the reference sequence (NC_058373.1: 115262322– 115282533, reverse complement) was used as the query in *blastn* searches against each database. The *blastn* were performed using the following parameters: *-max_target_seqs 5 - evalue 1 -word_size 7 -reward 1 -penalty -2 -gapopen 5 -gapextend 2*. The resulting alignments were processed using R packages (*seqinr* v.4.2-36 and *Biostrings* v.2.68.1), and manually inspected using AliView v.1.28 to verify sequence identity. The same approach was applied to investigate the reference NTCP sequence of lion (*Panthera leo*, XM_042943638.1), using whole-genome assemblies from four individuals (GCA_008795835.1, GCA_012313985.1, GCA_046254995.1, and GCA_046255005.1). The reference sequence (NC_056684.1: 114888058–114907759, reverse complement) was used as the query in *blastn* searches.

To investigate amino acid residues at aa 191 and 268 in lion NTCP, we also analyzed whole-genome sequencing reads derived from Sequence Read Archive (SRA) from 11 individuals (10 lions [*Panthera leo*] and 1 leopard [*Panthera pardus*] as a reference)[40]. Publicly available reads (SRR10764405–SRR10764415) were mapped to the reference sequence (NC_056684.1: 114888058–114907759, reverse complement) using BWA-MEM v.0.7.17, and amino acid residues were inferred from the aligned reads. Although the average genome coverage was reported to be relatively low (∼3.8x per individual)[40], and the number of reads mapped to the reference ranged from 0 to 21 per individual, this analysis allowed partial confirmation of the lion NTCP.

### Molecular dynamics simulations

We performed molecular dynamics (MD) simulations for four apo-like NTCP systems prepared by removal of preS1, the wild type human, cat, and lion NTCPs, and a lion mutant carrying the R191G/A268T substitution. The cryoEM structure of human NTCP-preS1 complex (PDB: 8HRY) was used as the initial structure for human NTCP, from which preS1 was removed. Stuctures for cat-NTCP, lion-NTCP, and lion-NTCP(R191G/A268T) in complex with preS1 were prepared by homology modeling using MODELLER[52] ver.9.25 with PDB: 8HRY as a template. Multiple sequence alignment among human, cat, and lion NTCPs performed by Clustal Omega[53] shows the sequence similarity of 84% between human-NTCP and cat-NTCP, 83% between human-NTCP and lion-NTCP, and 97% between cat-NTCP and lion-NTCP. The preS1-bound structure was used as a template to preserve the overall preS1-binding site architecture during homology modeling, and the apo-like initial structures were prepared by removing the preS1(2‒48) moiety from the complex structures. The membrane-water systems were prepared using Membrane Builder module implemented in CHARMM-GUI[54–56]. During setup in CHARMM-GUI, missing hydrogen atoms were added, according to the protonation states of residues determined by their pKa values at NTCP-preS1 complex estimated by PROPKA implemented in PDB 2PQR[57]: In human-NTCP, E257 was set to a protonated glutamic acid, and all the histidine residues except for H54 were protonated only N_δ_ atom (HSD in the charm force field), and H54 and were protonated only N_Ɛ_ atom (HSE in the charm force field). In the apo state of cat- and lion-NTCP systems, E257 and E296 were set to a protonated glutamic acid, and all the histidine residues were protonated only N_δ_ atom (HSD in the charm force field). In all the systems, the N- and C-termini in NTCP were set to NH_3_^+^ and COO^−^, respectively. In the lion-NTCP(R191G/A268T) mutant systems, the same protonation state for wild-type was employed. The R191G/A268T substitutions in lion-NTCP were performed in the setup process. The apo NTCP was embedded in a 70 Å x 70 Å 1-palmitoyl-2-oleoyl-glycero-3 -phosphocholin (POPC) bilayer in the x-y plane in the center of the rectangular unit cell. The orientation of NTCP relative to the membrane was set in the same as that of NTCP deposited in the Orientations of Proteins in Membranes database (PDB: 7FCI). After filling the unit cell with water molecules (TIP3water model), sodium and chloride counterions (NaCl) were added with 150 mM ion concentration.

All-atom conventional MD simulations were performed and analyzed by using the MD program package GROMACS ver. 2021.4[58] under periodic boundary conditions. The CHARMM36m force field[59–61], which is often used for membrane-protein systems, was used. Energy minimization and equilibration runs were performed according to the default setting in Membrane Builder; detailed conditions were the same as those of our previous simulations[62]. Then, three independent 1-µs production runs with different initial velocities and initial configuration were performed; each independent MD was performed from the energy minimization process. The NPT ensemble was used, and temperature and pressure were set at 300 K and 1 atm, respectively. The thermostat and barostat used were the Nose-Hoover scheme and the Parrinello–Rahman approach, respectively. The type of pressure coupling was semi-isotropic. The particle mesh Ewald method was used for evaluating electrostatic interactions, and the van der Waals interactions were smoothly truncated using the switching function within 10-12 Å range. Bond length involving hydrogen atoms was constrained by the P-LINCS algorithm, and a 2-fs timestep was used. MD simulations for the four systems were performed using the same procedure and conditions as described above. Snapshots were saved every 1 ns from a 1-µs MD simulation and used for trajectory analyses. Principal component analysis (PCA) of the NTCP structures was performed using C_α_ atoms for the 21-310 residues of NTCP after a structural alignment of the snapshots using the C_α_ atoms of 21-310 residues, and the PC axes were determined using three MD trajectories for Cat and Lion systems and the four cryo-EM structures (PDB: 7FCI, 8HRY, 7ZYI, and 7PQG). Contributions of PC1 and PC2 against the total degree of freedom were 28.4 % and 17.1%, respectively. Snapshots were projected onto the PC axes. The free energy landscapes shown as color maps (Fig. 6A and D) were evaluated by ‒ln*P*, where *P* is the existence probability calculated on the normalized histogram of 1 Å x 1 Å pixels on the PC1-PC2 surface.

### Statistics

Bar graphs show the means ± standard deviation with invidual data set, and experiments were repeated in triplicate. One-way or Two-way ANOVA were performed by GraphPad prism10.

## Supporting information

**Table S1. List of NTCP derived from mammalian species with GenBank ID.**

**S1 Fig. Immunofluorescence detection of preS1 binding to NTCPs derived from 23 animals.** In the preS1 binding assay performed as shown in Fig. 1, TAMRA-labelled preS1 (red), NTCP (green), and the nucleus (blue) were detected by immunefluorescence analysis. Merged patterns of the fluorescences are also shown. Myrcludex-B (myrB) was used as a positive control that inhibits preS1 binding to NTCPs. Scale bars, 100 µm.

**S2 Fig. Physiological functions of feline family-derived NTCPs. (A)** Subcellular localization of NTCPs derived from the indicated species as well as cadherin [as a plasma membrane (PM) marker] and the nucleus were examined by immunofluorescence analysis with a confocal microscopy, using the cells prepared as shown in Fig. 1B. Green, NTCP; red, cadherin, blue, nucleus. Merged patterns are also shown. **(B)** Transporter kinetics was examined in the cells expressing human, cat, and lion-NTCP by treatment with different concentrations of taurocholic acid (3.75, 7.5, 15, 30, 60, 120, 240 µM). *K_m_* and *V_max_* were calculated with GraphPad prism10. *K_m_* is 10.26, 27.66, 33.08 for human, cat, and lion NTCP, whereas *V_max_* normalized by the cell surface NTCP expression level is 5327, 6869, and 922 for human, cat, and lion NTCP, respectively.

**S3 Fig. Sequence analysis of NTCP derived from cat and lion. (A, B)** Multiple sequence alignment of the experimentally identified cat NTCP cDNA sequence with predicted and reference sequences from the database. Six cat NTCP sequences were aligned, one determined experimentally from the sequence of NTCP cDNA from a domestic cat liver [cDNA (from liver)], four predicted by whole-genome assemblies from public database (GCA_000003115.1, GCA_000181335.6, GCA_013340865.2, and GCA_016509815.2), and the reference sequence registered as mRNA in the NCBI RefSeq database (XM_003987782.5). Conserved amino acids are shown by dot. The visualization was generated using BioEdit Sequence Alignment Editor. DNA (a) and amino acid (b) sequences are shown. **(C, D)** Multiple sequence alignment of the predicted lion NTCP sequences and reference from the database. A total of five lion NTCP sequences were aligned, four predicted by whole-genome assemblies from public database (GCA_008795835.1, GCA_012313985.1, GCA_046254995.1, and GCA_046255005.1), and the reference sequence registered as mRNA in the NCBI RefSeq database (XM_042943638.1). DNA (c) and amino acid (d) sequences are shown. **(E)** Amino acid residues and the corresponding codons at aa 191 and 268 of lion NTCP sequences inferred from whole-genome sequencing reads of individual animals. Each tile represents the result for each position in one individual (SRR ID). Colors indicate amino acid types based on reference sequences; yellow for lion-type, cyan for cat-type, and gray for undetermined residues (ND). Residue and codon were inferred from short-read alignments using BWA-MEM v.0.7.17. and compared with the reference sequence (NC_056684.1). Figure visualization was conducted using R 4.3.2 (R Foundation for statistical Computing, Vienna, Austria).

**S4 Fig. Schematic representation for principal component analysis (PCA) in molecular dynamics simulations.** PC1 (red) axis corresponds to motion related to differences between the inward-facing and the outward-facing NTCP structures, while PC2 (blue) axis corresponds to motion related to differences between the preS1-binding and bile acid-binding NTCP structures. PC2 also incorporates motion related to differences between the inward-facing and bile acid-binding NTCP structures.

**S1 Data.** Source data for all experiments are shown.

## Data Availability

All MD and associated analysis data have been submitted to the Biological Structure Model Archive (BSM-Arc) under BSM-ID BSM00105 (https://bsma.pdbj.org/entry/105).

S1 Table

## Acknowledgments

The HDV-expression plasmid, pSVLD3, was kindly provided by Dr. John Taylor at Fox Chase Cancer Center. K.W. was supported by Japan Agency for Medical Research and Development (AMED) under grand numbers JP26fk0310525, JP26fk0310533, JP26fk0310547, JP23ama121007, JP26ak0101271, JP26wm0325079, JP26jm0110034; JSPS/MEXT under KAKENHI grant numbers JP24K02290, JP25K21767, JP25K21782, JP25K02691; JST MIRAIprogram grant number JPMJMI22G1.

## Author Contributions

**Conceptualization:** Koichi Watashi.

**Data Curation:** Junki Mifune, Toru Ekimoto, Junko S Takeuchi, Miki X Kawano, Akatsuki Saito, Mitsunori Ikeguchi.

**Formal Analysis:** Junki Mifune, Toru Ekimoto, Junko S Takeuchi, Miki X Kawano, Akatsuki Saito, Mitsunori Ikeguchi.

**Funding Acquisition:** Koichi Watashi.

**Investigation:** Junki Mifune, Chisa Kobayashi, Misaki Kurata, Kaho Shionoya. Project Administration: Koichi Watashi.

**Resources:** Camille Sureau.

**Supervision:** Koichi Watashi.

**Validation:** Junki Mifune, Toru Ekimoto, Chisa Kobayashi, Misaki Kurata, Kaho Shionoya.

**Writing – Original Draft Preparation:** Junki Mifune, Toru Ekimoto, Junko S Takeuchi, Miki X Kawano, Akatsuki Saito, Koichi Watashi.

**Writing – Review & Editing:** Chisa Kobayashi, Misaki Kurata, Kaho Shionoya.

**Competing Interests**

We declare that none of the authors has financial interests related to this work.

