## Supplemental Data 1 for "Hepadnavirus tropism determined by diversified NTCP functionality in Feline species"

**Contents:**

**Table S1**

**Figure S1 – S4**

**Table S1. List of NTCP derived from mammalian species with GenBank ID.**

| Common name | Species | Accession (amino acid) | Accession (mRNA) |
|---|---|---|---|
| Leopard | *Panthera pardus* | XP_019306946.1 | XM_019451401.1 |
| Tiger | *Panthera tigris* | XP_007095203.2 | XM_007095141.3 |
| Cat | *Felis catus* | XP_003987831.1 | XM_003987782.5 |
| Lion | *Panthera leo* | XP_042799572.1 | XM_042943638.1 |
| American mink | *Neovison vison* | XP_044088226.1 | XM_044232291.1 |
| Ferret | *Mustela putorius furo* | XP_004739077.1 | XM_004739020.3 |
| Dog | *Canis lupus familiaris* | XP_025298474.1 | XM_025442689.3 |
| Tree shrew | *Tupaia chinensis* | XP_006171565.1 | XM_006171503.3 |
| Capuchin | *Tufted capuchin* | XP_032101672.1 | XM_032245781.1 |
| Common marmoset | *Callithrix jacchus* | XP_035116722.1 | XM_035260831.1 |
| Human | *Homo sapiens* | NP_003040.1 | NM_003049.4 |
| Sumatran orangutan | *Pongo abelii* | XP_002824936.1 | XM_002824890.2 |
| Chimpanzee | *Pan troglodytes* | XP_003824149.1 | XM_003824101.2 |
| Golden snub-nosed monkey | *Rhinopithecus roxellana* | XP_010358049.1 | XM_010359747.2 |
| Drill | *Mandrillus leucophaeus* | XP_011856290.1 | XM_012000900.1 |
| Macaque | *Macaca* | NP_001270252.1 | NM_001283323.1 |
| Cow | *Bos taurus* | NP_001039804.1 | NM_001046339.1 |
| Goat | *Capra hircus* | XP_005686078.2 | XM_005686021.2 |
| Sheep | *Ovis aries* | XP_027827755.1 | XM_027971954.2 |
| woodchuck | *Marmota monax* | XP_015346065.1 | XM_015490579.1 |
| Chinese hamster | *Cricetulus griseus* | XP_035301832.1 | XM_035445941.1 |
| Mouse | *Mus musculus* | NP_001171032.1 | NM_001177561.1 |
| Rat | *Rattus norvegicus* | NP_058743.1 | NM_017047.2 |

#### Supplementary Figure legends

**S1 Fig. Immunofluorescence detection of preS1 binding to NTCPs derived from 23 animals.**

In the preS1 binding assay performed as shown in Fig. 1, TAMRA-labelled preS1 (red), NTCP (green), and the nucleus (blue) were detected by immunefluorescence analysis. Merged patterns of the fluorescences are also shown. Myrcludex-B (myrB) was used as a positive control that inhibits preS1 binding to NTCPs. Scale bars, 50 μm.

**S2 Fig. Physiological functions of feline family-derived NTCPs.**

**(A)** Subcellular localization of NTCPs derived from the indicated species as well as cadherin [as a plasma membrane (PM) marker] and the nucleus were examined by immunofluorescence analysis with a confocal microscopy, using the cells prepared as shown in Fig. 1B. Green, NTCP; red, cadherin, blue, nucleus. Merged patterns are also shown. **(B)** Transporter kinetics was examined in the cells expressing human, cat, and lion-NTCP by treatment with different concentrations of taurocholic acid (3.75, 7.5, 15, 30, 60, 120, 240 μM). $K_m$ and $V_{max}$ were calculated with GraphPad prism10. $K_m$ is 10.26, 27.66, 33.08 for human, cat, and lion NTCP, whereas $V_{max}$ normalized by the cell surface NTCP expression level is 5327, 6869, and 922 for human, cat, and lion NTCP, respectively.

**S3 Fig. Sequence analysis of NTCP derived from cat and lion.**

**(A, B)** Multiple sequence alignment of the experimentally identified cat NTCP cDNA sequence with predicted and reference sequences from the database. Six cat NTCP sequences were aligned, one determined experimentally from the sequence of NTCP cDNA from a domestic cat liver [cDNA (from liver)], four predicted by whole-genome assemblies from public database (GCA_000003115.1, GCA_000181335.6, GCA_013340865.2, and GCA_016509815.2), and the reference sequence registered as mRNA in the NCBI RefSeq database (XM_003987782.5). Conserved amino acids are shown by dot. The visualization was generated using BioEdit Sequence Alignment Editor. DNA (a) and amino acid (b) sequences are shown. **(C, D)** Multiple sequence alignment of the predicted lion NTCP sequences and reference from the database. A total of five lion NTCP sequences were aligned, four predicted by whole-genome assemblies from public database (GCA_008795835.1, GCA_012313985.1, GCA_046254995.1, and GCA_046255005.1), and the reference sequence registered as mRNA in the NCBI RefSeq database (XM_042943638.1). DNA (c) and amino acid (d) sequences are shown. **(E)** Amino acid residues and the corresponding codons at aa 191 and 268 of lion NTCP sequences inferred from whole-genome sequencing reads of individual animals. Each tile represents the result for each position in one individual (SRR ID). Colors indicate amino acid types based on reference sequences; yellow for lion-type, cyan for cat-type, and gray for undetermined residues (ND). Residue and codon were inferred from short-read alignments using BWA-MEM v.0.7.17. and compared with the reference sequence (NC_056684.1). Figure visualization was conducted using R 4.3.2 (R Foundation for statistical Computing, Vienna, Austria).

**S4 Fig. Schematic representation for principal component analysis (PCA) in molecular dynamics simulations.**

PC1 (red) axis corresponds to motion related to differences between the inward-facing and the outward-facing NTCP structures, while PC2 (blue) axis corresponds to motion related to differences between the preS1-binding and bile acid-binding NTCP structures. PC2 also incorporates motion related to differences between the inward-facing and bile acid-binding NTCP structures.

**S1 Data.** Source data for all experiments are shown.

### Supplementary Table 1

| Comon name | Species | Accession (amino acid) | Accession (mRNA) |
|---|---|---|---|
| Leopard | *Panthera pardus* | XP_019306946.1 | XM_019451401.1 |
| Tiger | *Panthera tigris* | XP_007095203.2 | XM_007095141.3 |
| Cat | *Felis catus* | XP_003987831.1 | XM_003987782.5 |
| Lion | *Panthera leo* | XP_042799572.1 | XM_042943638.1 |
| American mink | *Neovison vison* | XP_044088226.1 | XM_044232291.1 |
| Ferret | *Mustela putorius furo* | XP_004739077.1 | XM_004739020.3 |
| Dog | *Canis lupus familiaris* | XP_025298474.1 | XM_025442689.3 |
| Tree shrew | *Tupaia chinensis* | XP_006171565.1 | XM_006171503.3 |
| Capuchin | *Tufted capuchin* | XP_032101672.1 | XM_032245781.1 |
| Common marmoset | *Callithrix jacchus* | XP_035116722.1 | XM_035260831.1 |
| Human | *Homo sapiens* | NP_003040.1 | NM_003049.4 |
| Sumatran orangutan | *Pongo abelii* | XP_002824936.1 | XM_002824890.2 |
| Chimpanzee | *Pan troglodytes* | XP_003824149.1 | XM_003824101.2 |
| Golden snub-nosed monkey | *Rhinopithecus roxellana* | XP_010358049.1 | XM_010359747.2 |
| Drill | *Mandrillus leucophaeus* | XP_011856290.1 | XM_012000900.1 |
| Macaque | *Macaca* | NP_001270252.1 | NM_001283323.1 |
| Cow | *Bos taurus* | NP_001039804.1 | NM_001046339.1 |
| Goat | *Capra hircus* | XP_005686078.2 | XM_005686021.2 |
| Sheep | *Ovis aries* | XP_027827755.1 | XM_027971954.2 |
| woodchuck | *Marmota monax* | XP_015346065.1 | XM_015490579.1 |
| Chinese hamster | *Cricetulus griseus* | XP_035301832.1 | XM_035445941.1 |
| Mouse | *Mus musculus* | NP_001171032.1 | NM_001177561.1 |
| Rat | *Rattus norvegicus* | NP_058743.1 | NM_017047.2 |

### Fig. S1

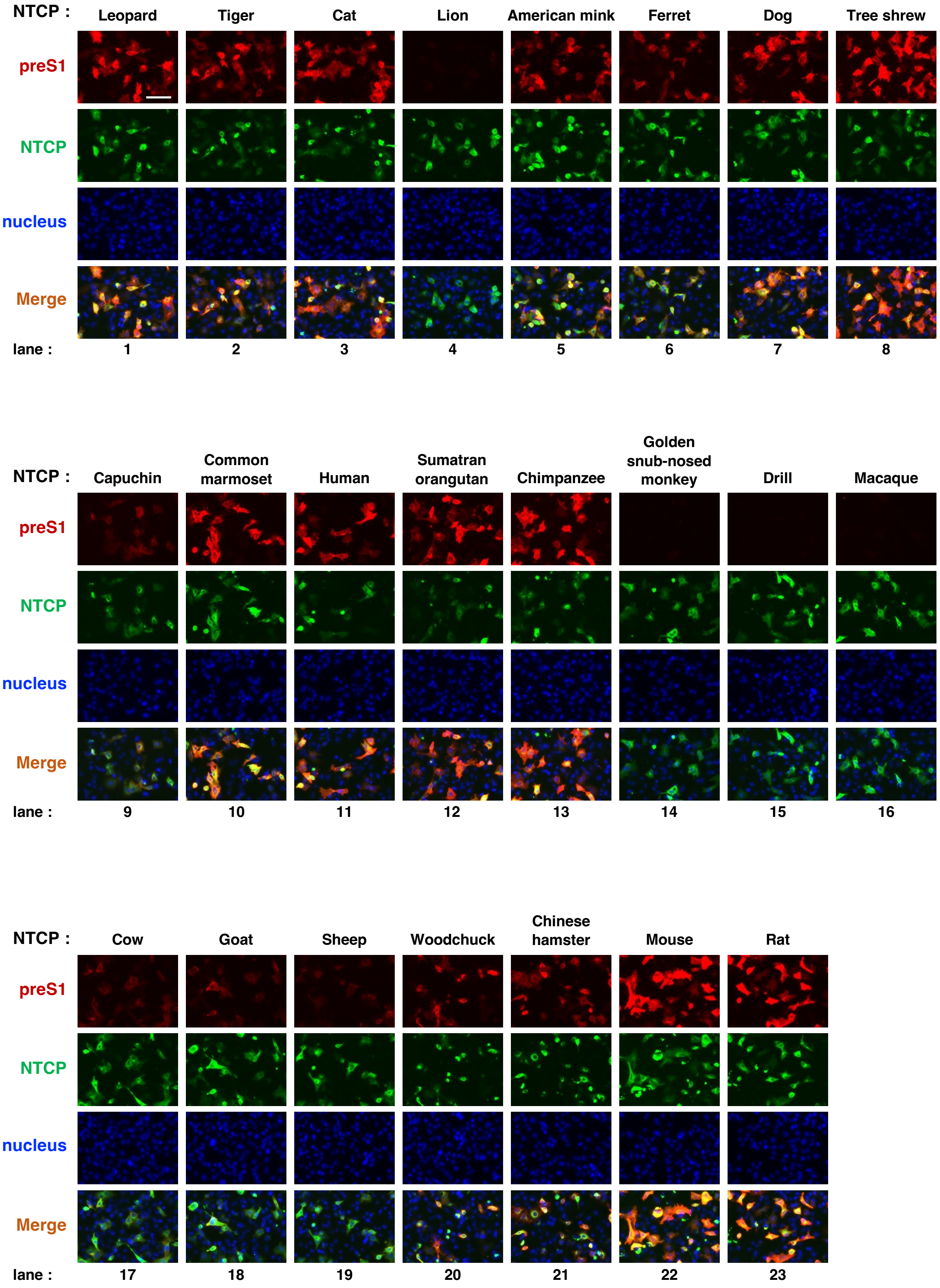
NTCP :
Leopard
Tiger
Cat
Lion
American mink
Ferret
Dog
Tree shrew
preS1
NTCP
nucleus
Merge
lane :
1
2
3
4
5
6
7
8
NTCP :
Capuchin
Common marmoset
Human
Sumatran orangutan
Chimpanzee
Golden snub-nosed monkey
Drill
Macaque
preS1
NTCP
nucleus
Merge
lane :
9
10
11
12
13
14
15
16
NTCP :
Cow
Goat
Sheep
Woodchuck
Chinese hamster
Mouse
Rat
preS1
NTCP
nucleus
Merge
lane :
17
18
19
20
21
22
23

#### Fig. S2

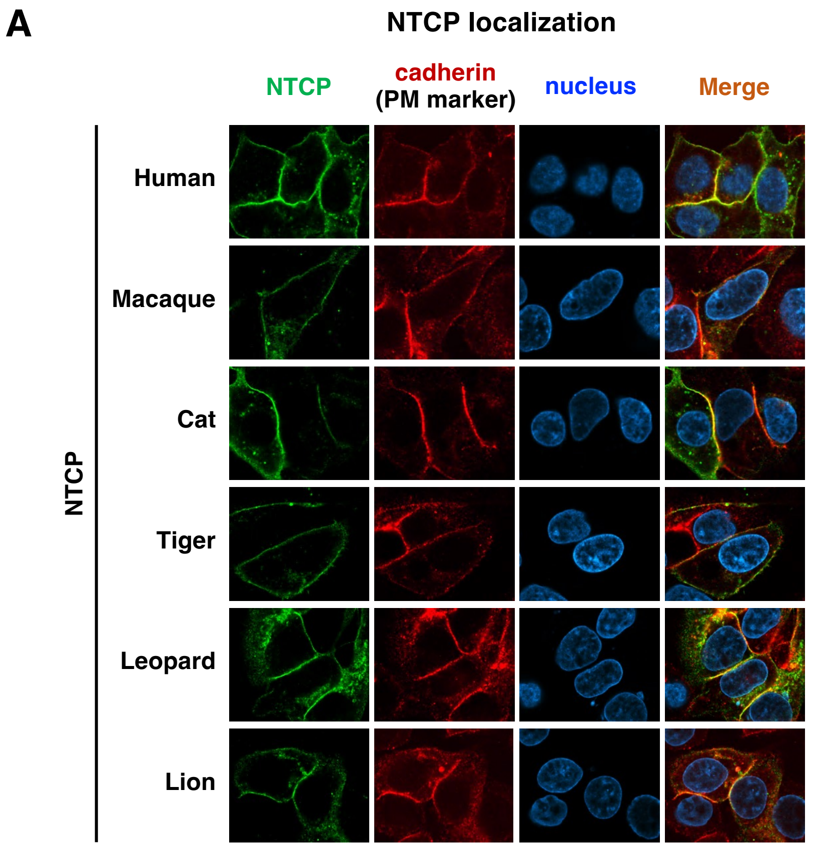

A
NTCP localization
NTCP
cadherin
(PM marker)
nucleus
Merge
NTCP
Human
Macaque
Cat
Tiger
Leopard
Lion

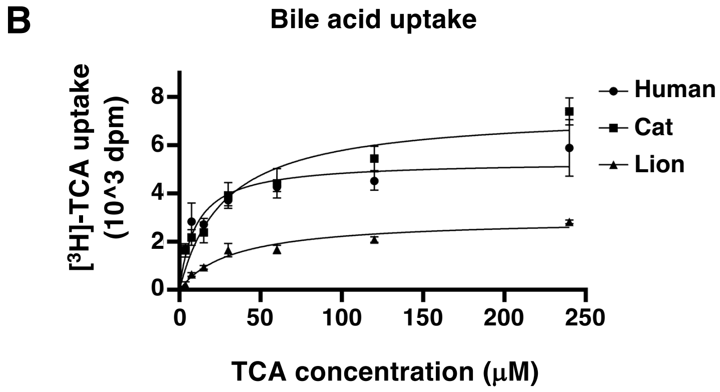

B
Bile acid uptake
[3H]-TCA uptake
(10^3 dpm)
0
2
4
6
8
0
50
100
150
200
250
TCA concentration (μM)
Human
Cat
Lion

### Fig. S3

#### A cat NTCP (DNA)

```
XM_003987782.5    ATGGAGCCCCACAATGTGACTGCCACGCTCAACTTCACCCTTCCACCCAACTTCGGCAAGCGCCCCACCGACAAGGCACTGAGCGTCATCCTGGTGTTCC 100
GCA_000003115.1   .................................................................................................... 100
GCA_000181335.6   .................................................................................................... 100
GCA_013340865.2   .................................................................................................... 100
GCA_016509815.2   .................................................................................................... 100
cDNA (from liver) .................................................................................................... 100

XM_003987782.5    TGCTCCTTATCATCATGCTCTCGCTGGGCTGCACCATGGAGTTCAGCAAGATCAGGGCTCACTTCTGGAAGCCTAAGGGGCTGGCCATCGCCCTGATAGC 200
GCA_000003115.1   .................................................................................................... 200
GCA_000181335.6   .................................................................................................... 200
GCA_013340865.2   .................................................................................................... 200
GCA_016509815.2   .................................................................................................... 200
cDNA (from liver) .................................................................................................... 200

XM_003987782.5    ACAGTACGGCATCATGCCCCTCACCGCCTTTGCACTGGGCAAGGTCTTCCAGCTCAACAACATTGAGGCGCTGGCCATCCTTGTCTGCGGCTGCTCACCC 300
GCA_000003115.1   .................................................................................................... 300
GCA_000181335.6   .................................................................................................... 300
GCA_013340865.2   .................................................................................................... 300
GCA_016509815.2   .................................................................................................... 300
cDNA (from liver) .................................................................................................... 300

XM_003987782.5    GGGGGGACCCTGTCCAACATCTTCAGTCTGGCCATGAAGGGGGACATGAACCTCAGTATCGTGATGACCACCTGCTCCACCTTCTGTGCCCTGGGCATGA 400
GCA_000003115.1   .................................................................................................... 400
GCA_000181335.6   .................................................................................................... 400
GCA_013340865.2   .................................................................................................... 400
GCA_016509815.2   .................................................................................................... 400
cDNA (from liver) .................................................................................................... 400

XM_003987782.5    TGCCCCTCCTCCTGTATGTCTACTCCAGGGGGATCTATTCCGGGGATCTGAAGGACAAGGTACCTTATGGAGGCATTGTGATATCACTAATCCTGGTTCT 500
GCA_000003115.1   .................................................................................................... 500
GCA_000181335.6   .................................................................................................... 500
GCA_013340865.2   .................................................................................................... 500
GCA_016509815.2   .................................................................................................... 500
cDNA (from liver) .................................................................................................... 500

XM_003987782.5    CATTCCCTGTGCCACTGGGATCTTCCTCAATGCCAAACGGCCACAATATGTACCCTATGTCAAGAAGGGAGGGACCATCACCATGCTTTTGTTAAGTGTG 600
GCA_000003115.1   .................................................................................................... 600
GCA_000181335.6   .................................................................................................... 600
GCA_013340865.2   .................................................................................................... 600
GCA_016509815.2   .................................................................................................... 600
cDNA (from liver) .................................................................................................... 600

XM_003987782.5    GCCATCATAGCGCTCTCGGTCATCAACGTGGGCAAGAGTATCATGTTTGTCATGACACCACACTTGCTGGCCACCTCCTCCCTGATGCCTTTCATCGGCT 700
GCA_000003115.1   .................................................................................................... 700
GCA_000181335.6   .................................................................................................... 700
GCA_013340865.2   .................................................................................................... 700
GCA_016509815.2   .................................................................................................... 700
cDNA (from liver) .................................................................................................... 700

XM_003987782.5    TCCTGCTGGGCTACATCCTCTCTGCTCTCTTCCGCCTCAATGGACGGTGCAGGCGCACTGTCAGCATGGAAACTGGATGCCAAAACGTTCAGCTTTGTTC 800
GCA_000003115.1   .................................................................................................... 800
GCA_000181335.6   .................................................................................................... 800
GCA_013340865.2   .................................................................................................... 800
GCA_016509815.2   ........................................C........................................................... 800

XM_003987782.5    ATCTCCATCTTTCGTTGCTATGAGAAAATGAAGCCTTCCAAAGATAAAACAAAAATGACCTACACAGCTGCTCCAACTGAAGACACGATTCCAGGAGCTC 1000
GCA_000003115.1   .................................................................................................... 1000
GCA_000181335.6   .................................................................................................... 1000
GCA_013340865.2   .................................................................................................... 1000
GCA_016509815.2   .................................................................................................... 1000
cDNA (from liver) .................................................................................................... 1000

XM_003987782.5    TAGGAAATGGCACCCACAAAGGGGAGGAATGCTCCCCTTGCACAGCCTAG 1050
GCA_000003115.1   .............A.................................... 1050
GCA_000181335.6   .............A.................................... 1050
GCA_013340865.2   .................................................. 1050
GCA_016509815.2   .................................................. 1050
cDNA (from liver) .................................................. 1050
```

#### B cat NTCP (amino acid)

```
XM_003987782.5    MEPHNVTATLNFTLPPNFGKRPTDKALSVILVFLLLIIMLSLGCTMEFSKIRAHFWKPKGLAIALIAQYGIMPLTAFALGKVFQLNNIEALAILVCGCSP 100
GCA_000003115.1   .................................................................................................... 100
GCA_000181335.6   .................................................................................................... 100
GCA_013340865.2   .................................................................................................... 100
GCA_016509815.2   .................................................................................................... 100
cDNA (from liver) .................................................................................................... 100

XM_003987782.5    GGTLSNIFSLAMKGDMNLSIVMTTCSTFCALGMMPLLLYVYSRGIYSGDLKDKVPYGGIVISLILVLIPCATGIFLNAKRPQYVPYVKKGGTITMLLLSV 200
GCA_000003115.1   .................................................................................................... 200
GCA_000181335.6   .................................................................................................... 200
GCA_013340865.2   .................................................................................................... 200
GCA_016509815.2   .................................................................................................... 200
cDNA (from liver) .................................................................................................... 200

XM_003987782.5    AIIALSVINVGKSIMFVMTPHLLATSSLMPFIGFLLGYILSALFRLNGRCRRTVSMETGCQNVQLCSTILNVTFPPQVIGPLFFFPLLYMIFQLGEGVLL 300
GCA_000003115.1   .................................................................................................... 300
GCA_000181335.6   .................................................................................................... 300
GCA_013340865.2   .................................................................................................... 300
GCA_016509815.2   .................................................................................................... 300
cDNA (from liver) .................................................................................................... 300

XM_003987782.5    ISIFRCYEKMKPSKDKTKMTYTAAPTEDTIPGALGNGTHKGEECSPCTA* 350
GCA_000003115.1   .................................................* 350
GCA_000181335.6   .................................................* 350
GCA_013340865.2   .................................................* 350
GCA_016509815.2   .................................................* 350
cDNA (from liver) .................................................* 350
```

**Fig. S3** (continued)

**C** lion Ntcp (DNA)

```
XM_042943638.1   ATGGAGCCCCACAATGTGACTGCCACGCTCAACTTCACCCTCCCACCCAACTTCGGCAAGCGCCCCACCGACAAGGCACTGAGCGTCATCCTGGTGTTCC 100
GCA_008795835.1  .................................................................................................... 100
GCA_012313985.1  .................................................................................................... 100
GCA_046254995.1  .....................................................T.............................................. 100
GCA_046255005.1  .................................................................................................... 100

XM_042943638.1   TGCTCCTTATCATCATGCTCTCACTGGGCTGCACCATGGAGTTCAGCAAGATCAAGGCTCACTTCTGGAAGCCTAAGGGGCTGGCCATCGCCCTGATAGC 200
GCA_008795835.1  ......................G............................................................................. 200
GCA_012313985.1  .................................................................................................... 200
GCA_046254995.1  ......................G............................................................................. 200
GCA_046255005.1  ......................G............................................................................. 200

XM_042943638.1   ACAGTACGGCATCATGCCCCTCACCGCCTTTGCACTGGGCAAGGTCTTCCAGCTCAACAACATTGAGGCGCTGGCCATCCTTGTCTGCGGCTGCTCACCC 300
GCA_008795835.1  .................................................................................................... 300
GCA_012313985.1  .................................................................................................... 300
GCA_046254995.1  .................................................................................................... 300
GCA_046255005.1  .................................................................................................... 300

XM_042943638.1   GGGGGGACCCTGTCCAACATCTTCAGTCTGGCCATGAAGGGGGACATGAACCTCAGTATCATGATGACCACCTGCTCCACCTTCTGTGCCCTGGGCATGA 400
GCA_008795835.1  .................................................................................................... 400
GCA_012313985.1  .................................................................................................... 400
GCA_046254995.1  .................................................................................................... 400
GCA_046255005.1  .................................................................................................... 400

XM_042943638.1   TGCCCCTCCTCCTGTATGTCTACTCCAGGGGGATCTATGCCGGGGATCTGAAGGACAAGGTACCTTATGGAGGCATTGTGATATCACTAATCCTGGTTCT 500
GCA_008795835.1  .................................................................................................... 500
GCA_012313985.1  .................................................................................................... 500
GCA_046254995.1  .................................................................................................... 500
GCA_046255005.1  .................................................................................................... 500

XM_042943638.1   CATTCCCTGTGCCACTGGGATCTTCCTCAATGCCAAACGGCCACAATATGTACCCTATGTCAAGAAGGGAAGGACCATCACCACGCTTTTGTTAAGTGTG 600
GCA_008795835.1  .................................................................................................... 600
GCA_012313985.1  .................................................................................................... 600
GCA_046254995.1  .................................................................................................... 600
GCA_046255005.1  .................................................................................................... 600

XM_042943638.1   GCCATCATAGCGCTCTCAGTCATCAATGTGGGCAAGAGTATCATGCTTGTCATGACCCCACACTTGCTGGCCACCTCCTCCCTGATGCCTTTCATCGGCT 700
GCA_008795835.1  .................................................................................................... 700
GCA_012313985.1  .................................................................................................... 700
GCA_046254995.1  .................................................................................................... 700
GCA_046255005.1  .................................................................................................... 700

XM_042943638.1   TCCTGTTGGGCTACATCCTCTCTGCTCTCTTCCACCTCAATGGACGGTGCAGGCGCACTGTCAGCATGGAAACTGGATGCCAAAACGTTCAACTTTGTTC 800
GCA_008795835.1  .................................................................................................... 800
GCA_012313985.1  .................................................................................................... 800
GCA_046254995.1  .................................................................................................... 800
GCA_046255005.1  .................................................................................................... 800

XM_042943638.1   TGCCATTCTCAATGTGACGTTCCCCCCTCAAGTCATTGGGCCACTTTTATTCTTCCCTCTCCTCTACATGATTTTCCAGCTTGGAGAAGGGGTTCTTCTC 900
GCA_008795835.1  .................................................................................................... 900
GCA_012313985.1  .................................................................................................... 900
GCA_046254995.1  .................................................................................................... 900
GCA_046255005.1  .................................................................................................... 900

XM_042943638.1   ATCTCCATCTTTCGTTGCTATGAGAAAATCAAGCCTTCCAAAGATAAAACAAAAATGACCTACACAGCTGCTCCAACTGAAGACACGATTCCAGGAGCTC 1000
GCA_008795835.1  .................................................................................................... 1000
GCA_012313985.1  .................................................................................................... 1000
GCA_046254995.1  .................................................................................................... 1000
GCA_046255005.1  .................................................................................................... 1000

XM_042943638.1   TAGGAAATGGCACCCACAAAGGGGAGGAATGCTCCCCTTGCACAGCCTAG 1050
GCA_008795835.1  .................................................. 1050
GCA_012313985.1  .................................................. 1050
GCA_046254995.1  .................................................. 1050
GCA_046255005.1  .................................................. 1050
```

**D** lion Ntcp (amino acid)

```
XM_042943638.1   MEPHNVTATLNFTLPPNFGKRPTDKALSVILVFLLLIIMLSLGCTMEFSKIKAHFWKPKGLAIALIAQYGIMPLTAFALGKVFQLNNIEALAILVCGCSP 100
GCA_008795835.1  .................................................................................................... 100
GCA_012313985.1  .................................................................................................... 100
GCA_046254995.1  .................................................................................................... 100
GCA_046255005.1  .................................................................................................... 100

XM_042943638.1   GGTLSNIFSLAMKGDMNLSIMMTTCSTFCALGMMPLLLYVYSRGIYAGDLKDKVPYGGIVISLILVLIPCATGIFLNAKRPQYVPYVKKGRTITTLLLSV 200
GCA_008795835.1  .................................................................................................... 200
GCA_012313985.1  .................................................................................................... 200
GCA_046254995.1  .................................................................................................... 200
GCA_046255005.1  .................................................................................................... 200

XM_042943638.1   AIIALSVINVGKSIMLVMTPHLLATSSLMPFIGFLLGYILSALFHLNGRCRRTVSMETGCQNVQLCSAILNVTFPPQVIGPLLFFPLLYMIFQLGEGVLL 300
GCA_008795835.1  .................................................................................................... 300
GCA_012313985.1  .................................................................................................... 300
GCA_046254995.1  .................................................................................................... 300
GCA_046255005.1  .................................................................................................... 300

XM_042943638.1   ISIFRCYEKIKPSKDKTKMTYTAAPTEDTIPGALGNGTHKGEECSPCTA* 350
GCA_008795835.1  .................................................* 350
GCA_012313985.1  .................................................* 350
GCA_046254995.1  .................................................* 350
GCA_046255005.1  .................................................* 350
```

#### Fig. S3 (continued)

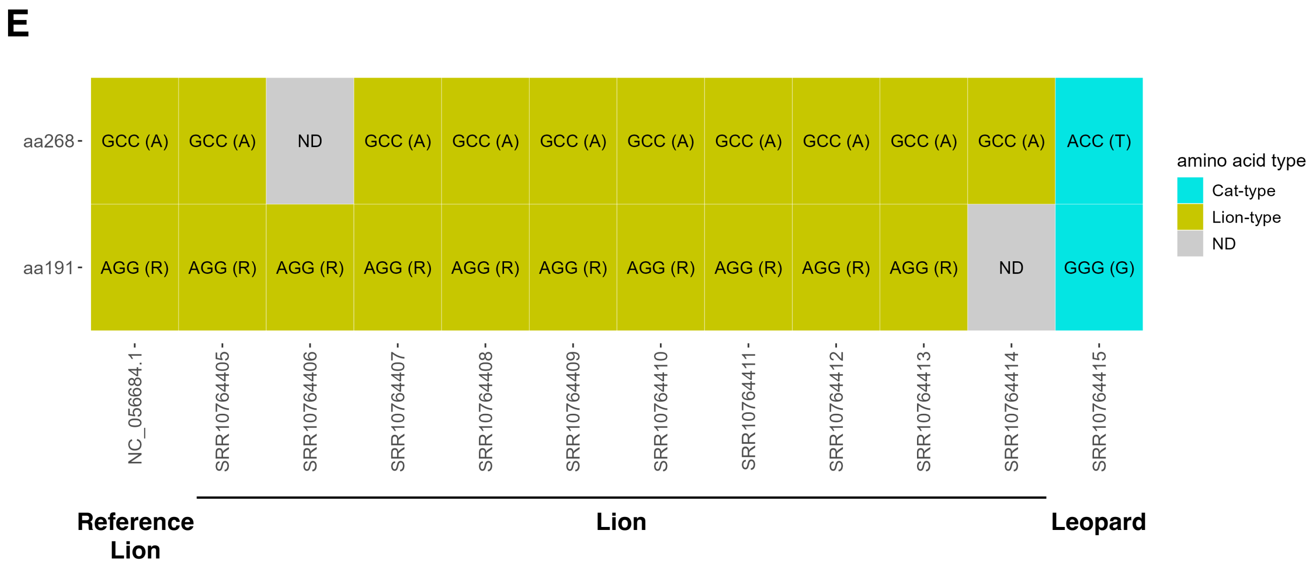

### Fig. S4

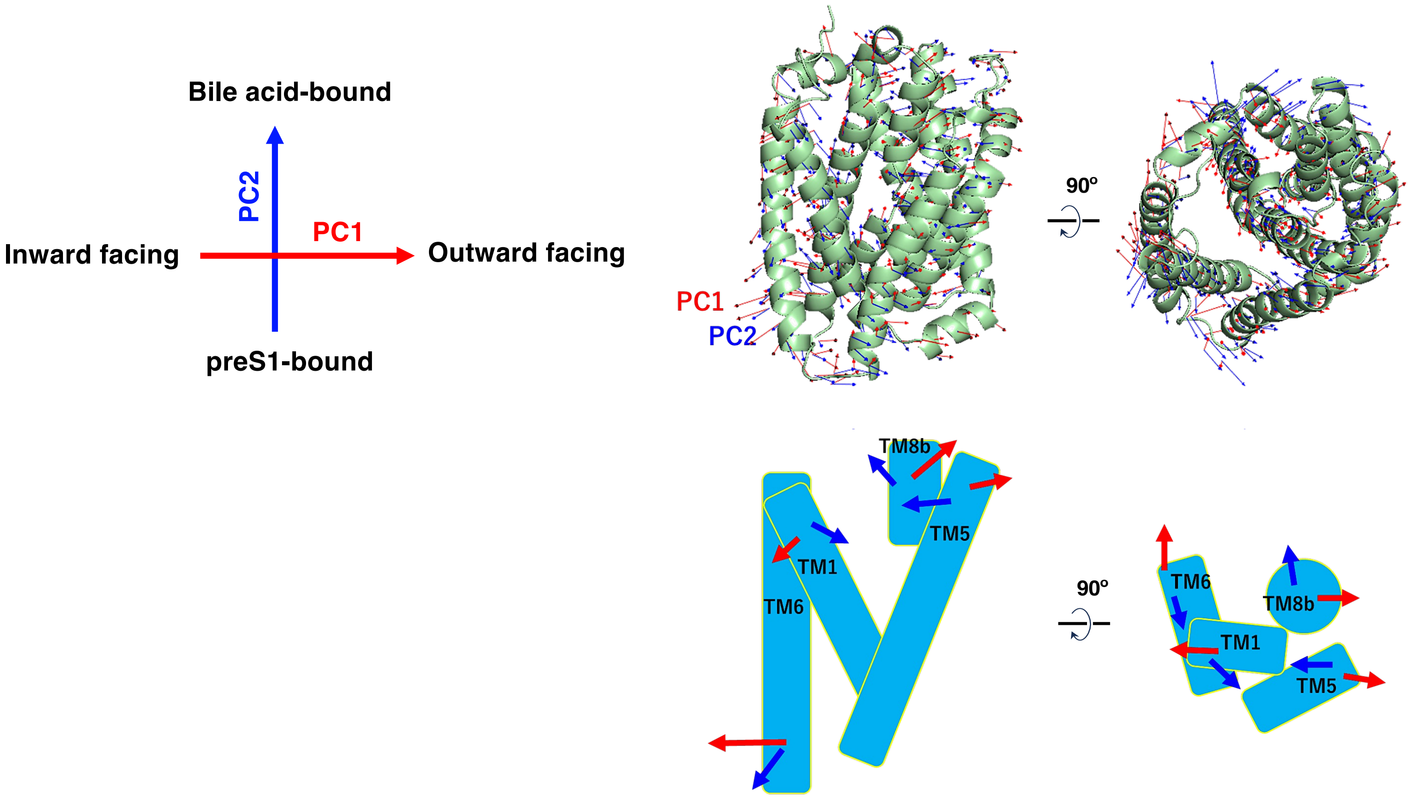

Bile acid-bound
PC2
PC1
Inward facing
Outward facing
preS1-bound
PC1
PC2
90º
TM8b
TM5
TM1
TM6
90º
TM6
TM8b
TM1
TM5
